# Dynamic Ry_sto_ receptor remodeling controls its ability to confer extreme resistance

**DOI:** 10.64898/2025.12.30.696988

**Authors:** Marta Grech-Baran, Juan Camilo Ochoa, Teresa Vargas-Cortez, Małgorzata Lichocka, Izabela Barymow-Filoniuk, J. T. Poznański, Magdalena Krzymowska

## Abstract

Plant nucleotide-binding leucine-rich repeat receptors (NLRs) mediate effector-triggered immunity (ETI), often accompanied by a hypersensitive response (HR). Conversely, extreme resistance (ER) provides exceptionally rapid and effective antiviral protection without visible cell death, yet the molecular and physiological mechanisms underlying ER remain poorly defined. The TIR-NLR receptor Ry_sto_ recognizes the coat protein (CP) of potato virus Y (PVY) and can trigger either ER or HR depending on the context. Here, we demonstrate that the efficacy of Ry_sto_-mediated ER relies on the orchestration of various defense mechanisms. These involve transcriptional priming, and spatial redistribution of the receptor and a subset of chaperones. Ry_sto_-expressing lines exhibit a preactivated immune state, including increased expression of genes encoding glycine-rich proteins associated with the cell wall interface and thickness. Upon PVY infection, *Ry_sto_* plants undergo rapid transcriptome reprogramming, including redox- and defense-related pathways. At the protein level, Ry_sto_ initially forms a pre-active complex with Hsp70 and CP-interacting cochaperonins (CPIPs) which is remodeled upon PVY CP binding. PVY CP competes for Hsp70, enabling Ry_sto_ to oligomerize into an active resistosome. This process is further accompanied by relocalization of a subpool of the receptor toward an interface of plasma membrane, cell wall. Our research reveals a chaperone-mediated activation mechanism and spatial immune repositioning that differentiate ER responses from traditional HR, offering a detailed system for achieving durable antiviral resistance in plants.

## Introduction

Plants rely on a multilayered immune system to perceive invading pathogens and deploy appropriate defenses. Pattern-triggered immunity (PTI) is initiated via detection of conserved pathogen-associated molecular patterns at the cell surface, while effector-triggered immunity (ETI) is mediated by intracellular nucleotide-binding leucine-rich repeat receptors (NLRs) (Ngou et al., 2022). NLRs contain an N-terminal signaling domain (CC or TIR), a central NB-ARC nucleotide-binding module, and C-terminal leucine-rich repeats that contribute to pathogen effector recognition (Jones et al., 2024). Upon effector perception, many NLRs oligomerize into resistosome complexes that activate robust defense signaling (Chai et., 2023; Jones et al., 2024).

A frequent outcome of ETI is the hypersensitive response (HR), a localized programmed cell death that limits pathogen growth (Coll et al., 2011) However, some NLRs trigger an even more robust antiviral defense known as extreme resistance (ER), which prevents detectable viral replication without inducing visible cell death (Bendahmane et al., 1994). ER is among the most effective forms of resistance in plants, yet its molecular and physiological underpinnings remain poorly understood. Only a few NLRs — including *Rx*, *PVR4*, *Rsv* loci (Bendahmane et al., 1994; Kim et al., 2015; Maroof et al., 2008), and the TIR-NLR *Ry_sto_* — are known to confer ER, and the mechanisms that distinguish ER from HR are largely unresolved.

*Ry_sto_* recognizes the coat protein (CP) of potato virus Y (PVY), a potyvirus of significant economic importance affecting potatoes (Grech-Baran et al., 2020). Depending on genetic and environmental factors, *Ry_sto_* activation can result in either ER or HR, although the mechanisms underlying these outcomes remain unclear. Previous research confirmed that Ry_sto_ is sufficient for PVY perception and ER induction in transgenic potato, with its ER phenotype being notably broad and strong across multiple PVY strains (Grech-Baran et al., 2022).

Global transcriptome analyses in various potato cultivars have shown rapid transcriptional changes upon PVY infection, with resistant genotypes exhibiting early modifications in stress, metabolic, and defense-related genes (ref). In contrast, susceptible plants tend to delay or display divergent gene expression patterns (Goyer et al., 2015). In Payette Russet, which carries *Ry_sto_*, transcriptome profiling at 24 hours post-inoculation identified extensive shifts in metabolic and defense pathways, implying that primary immune responses may occur earlier than previously sampled (Ross et al., 2022). Early differences in defense gene induction have been observed between resistant and susceptible cultivars within 4–12 hours post-infection (Beabler et al., 2009). Despite highlighting rapid immune responses, the variation in early sampling times and the use of different PVY strains and genetic backgrounds make it difficult to link transcriptomic signatures to Ry_sto_-mediated ER.

Successful virus infection requires host factors, including molecular chaperones. PVY exploits Hsp70 and Hsp40/DnaJ cochaperones to assemble movement-competent ribonucleoprotein complexes at cell walls and plasmodesmata, facilitating intercellular spread (Hoffius et al., 2007; Hafren et al., 2010). PVY CP interacts with Hsp70 and CP-interacting cochaperonins (CPIP1 and CPIP2a) early during infection, raising the possibility that these host pathways might also contribute to immune activation. In this study, we combine early-time-point transcriptomics with cell biological and biochemical analyses to define how Ry_sto_ integrates virus recognition with downstream defense activation. We show that Ry_sto_ expression confers a constitutively primed immune state characterized by elevated defense gene expression and glycine-rich protein (GRP) transcripts. Virus inoculation triggers rapid and extensive transcriptome reprogramming in *Ry_sto_* plants, including activation of redox and secondary metabolic pathways. At the cellular level, upon infection Ry_sto_ relocalizes toward the interface of plasma membrane and cell wall and is detectable in the apoplast, suggesting that spatial repositioning contributes to early virus perception and possibly inhibition of its spreading. At the molecular level, Ry_sto_ dimers form a pre-active complex with Hsp70 and CPIPs that is remodeled upon PVY CP engagement, enabling Ry_sto_ oligomerization into an active resistosome.

This integrated model highlights important physiological and mechanistic features that distinguish extreme resistance apart from classic HR.

## Results

### Potato plants expressing Ry_sto_ show rapid changes in the transcriptomic profile in response to PVY

To investigate the early molecular events underlying Ry_sto_-mediated extreme resistance (ER), we analyzed transcriptomic changes following its introduction into a PVY-susceptible potato background. We selected the cultivar Russet Burbank, which is highly susceptible to PVY, and generated transgenic lines expressing *Ry_sto_*. Resistance of these lines to PVY was previously confirmed (Grech-Baran et al., 2020). For transcriptome analysis, leaves were collected from a selected homogeneous *Ry_sto_*-Russet line and from wild-type (WT) Russet plants at three time points: prior to inoculation, immediately after inoculation (0 h), and 3 h post-inoculation (3 h). Mock-inoculated samples were included for each genotype and time point as controls.

To evaluate viral load in each sample, the reads were mapped to the PVY genome, revealing similar viral loads across all virus-inoculated plants, regardless of genotype (Table S1). Subsequently, Principal Components Analysis (PCA) was performed on the normalized read counts mapped to the *S. tuberosum* genome to evaluate the experiment’s quality and facilitate rapid visualization of the effects of each factor considered. The obtained results revealed that samples taken before inoculation or immediately afterward clustered together regardless of treatment, whereas the 3 h post-inoculation samples separated clearly by both treatment (PVY vs mock) and genotype (*Ry_sto_* vs WT) (Fig. 1A). Notably, *Ry_sto_* and WT plants also segregated by genotype prior to infection, indicating that *Ry_sto_* expression establishes a distinct basal transcriptomic state reminiscent of immune priming (Figure 1a).

**Figure 1.**
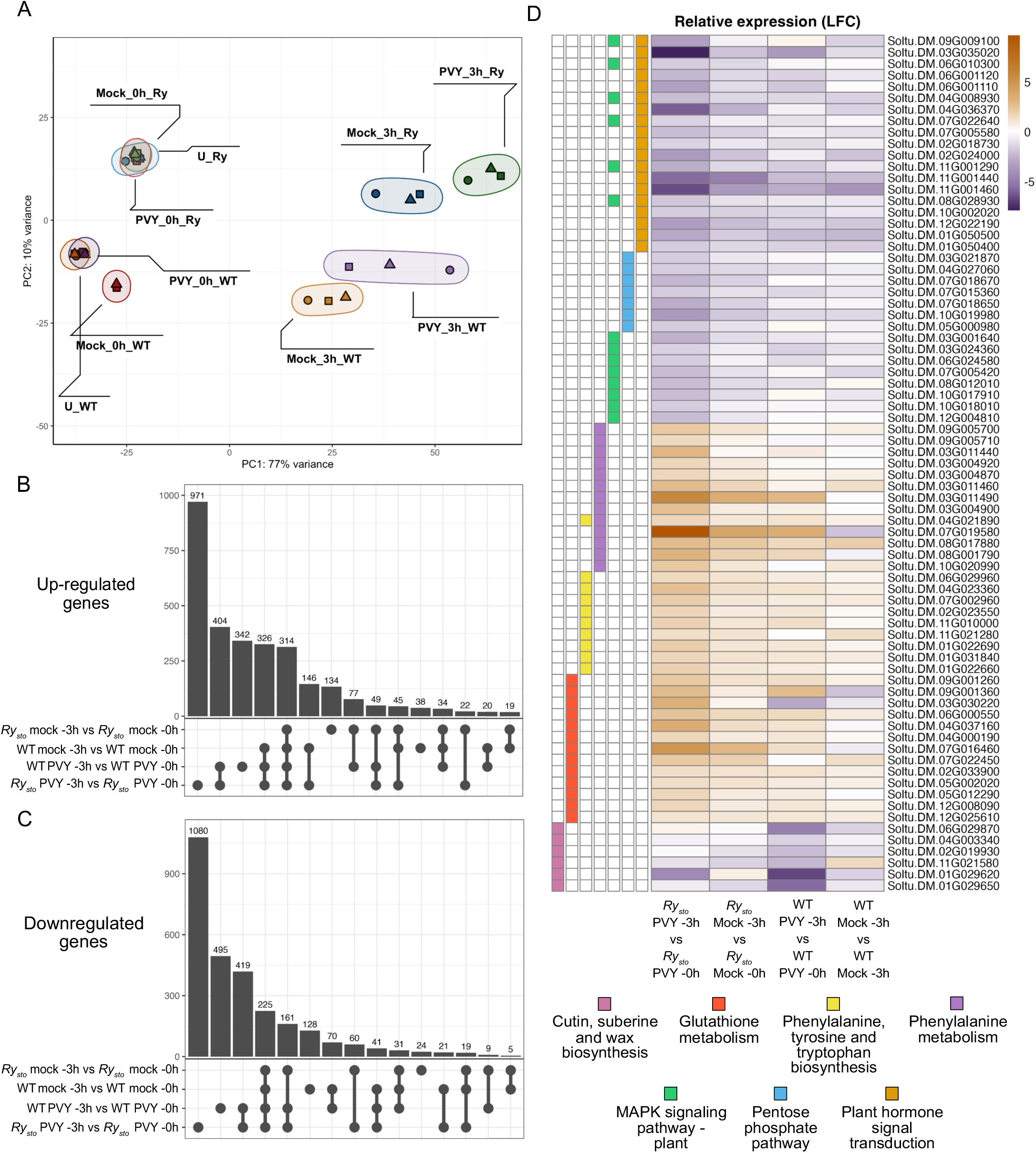
*Ry_sto_* plants exhibit a rapid and distinctive transcriptomic response to PVY inoculation. (A) Principal component analysis (PCA) of variance-stabilized gene expression counts across all samples. The first two components (PC1 and PC2) explain 77% and 10% of the total variance, respectively. Each point represents a biological replicate, with shapes indicating replicates and colors denoting genotype, treatment, and time-point combinations. (B, C) Upset plots showing the overlap of up-regulated (B) and down-regulated (C) genes between 0 h and 3 h post-inoculation. Vertical bars indicate the number of genes in each intersection, as shown by connected black dots. Differentially expressed genes were defined by |log2 fold-change| > 1 and adjusted p-value < 0.05. (D). Heatmap of log2 fold-changes for comparisons between 0 h and 3 h post-inoculation. Genes were selected from KEGG categories that were significantly enriched (q-value < 0.1).

Subsequently, we performed differential gene expression analysis with DESeq2 to identify genes and gene sets involved in the early response to PVY. The highest number of Differentially Expressed Genes (DEGs) was revealed in comparisons between the 3 h and 0 h time points after PVY inoculation in both *Ry_sto_* and WT plants, using a threshold of | log2 (Fold Change) | > 1 and a *p*-adjusted value < 0.05 (Table S1). Next, we compared the obtained DEG sets to identify genes differentially expressed exclusively in either *Ry_sto_* or WT plants 3 h after PVY inoculation. In total, we identified 971 up-regulated and 1,080 down-regulated genes only in *Ry_sto_*, and 342 up-regulated and 495 down-regulated genes only in WT plants (Figures 1B and 1C).

Enrichment analysis using Kyoto Encyclopedia of Genes and Genomes (KEGG) categories with the clusterProfiler R package, considering pathways significant at an FDR-adjusted p < 0.05, identified six significantly enriched pathways in *Ry_sto_* plants: three from up-regulated genes and three from down-regulated genes. The up-regulated genes in the *Ry_sto_* genotype were enriched in glutathione metabolism, phenylalanine metabolism, and phenylalanine, tyrosine, and tryptophan biosynthesis. This pattern indicates a potential activation of defense mechanisms involving antioxidants, antimicrobial compounds, pathogenesis-related (PR) proteins, cell wall lignification. In contrast, downregulated categories included MAPK signaling, pentose phosphate, and plant hormone pathways, particularly auxin signaling, consistent with a shift toward a defense-oriented, low-growth state. A unique enriched pathway identified in WT plants was related to cutine, suberin, and wax biosynthesis, suggesting a specifc response aimed at fortification of the cell wall (Fig.1C).

Overall, these results show that transcriptional program of *Ry_sto_* plants both makes them resilient to pathogens and allows for a rapid defense, while susceptible plants exhibit a weaker and/or delayed response.

### *Ry_sto_* plants display constitutive defense signatures and elevated glycine-rich protein expression

Since we detected distinctive constitutive transcriptomic signatures in *Ry_sto_* compared to WT plants, we decided to perform direct comparisons of mock-inoculated, PVY-inoculated, and untreated plants. We obtained a shared set of 62 genes with higher expression and 43 with lower expression in the transgenic plants (Figure 2A, Supp. dataset 1). Of the 62 genes upregulated in *Ry_sto_* plants 12 encode pathogenesis-related (PR) proteins—specifically Bet v I family members—along with two WRKY transcription factors and other immune-related genes, including flavin monooxygenases, TGA transcription factors, and a resistance gene homolog. In contrast, down-regulated genes were enriched in photosynthetic functions, such as chlorophyll-binding proteins (Supp. dataset 2). These findings suggest a constitutive activation of defense responses that likely occurs prior to viral colonization. Among constitutively upregulated genes, a cluster of glycine-rich protein (GRP)-encoding genes on chromosome 9 was prominent with the identifiers *Soltu.DM.09G030400, Soltu.DM.09G030390* and *Soltu.DM.09G030380* (Fig. 2B, 2C). Two additional orthologs, *Soltu.DM.09G030370* and *Soltu.DM.09G030410,* were identified using Orthofinder and showed higher and lower expression in *Ry_sto_* plants, respectively. Further sequence analysis of the candidate GRPs revealed a secretion peptide (SP) at their N-termini, followed by the glycine-rich region (GRR). With the exception of Soltu.DM.09G030400, the remaining four proteins likely belong to class V of GRPs, since they contain mixed patterns of (GGX)n/(GXGX)n repeats and a relatively high glycine content (Table S2) (Czolpinska & Rurek, 2018). On the other hand, Soltu.DM.09G030400 has a comparatively lower percentage of glycine residues and (GGX)n/(GXGX)n repeats but exhibits the highest baseline expression among all the *GRP* genes in the locus studied (Fig. 2C).

**Figure 2.**
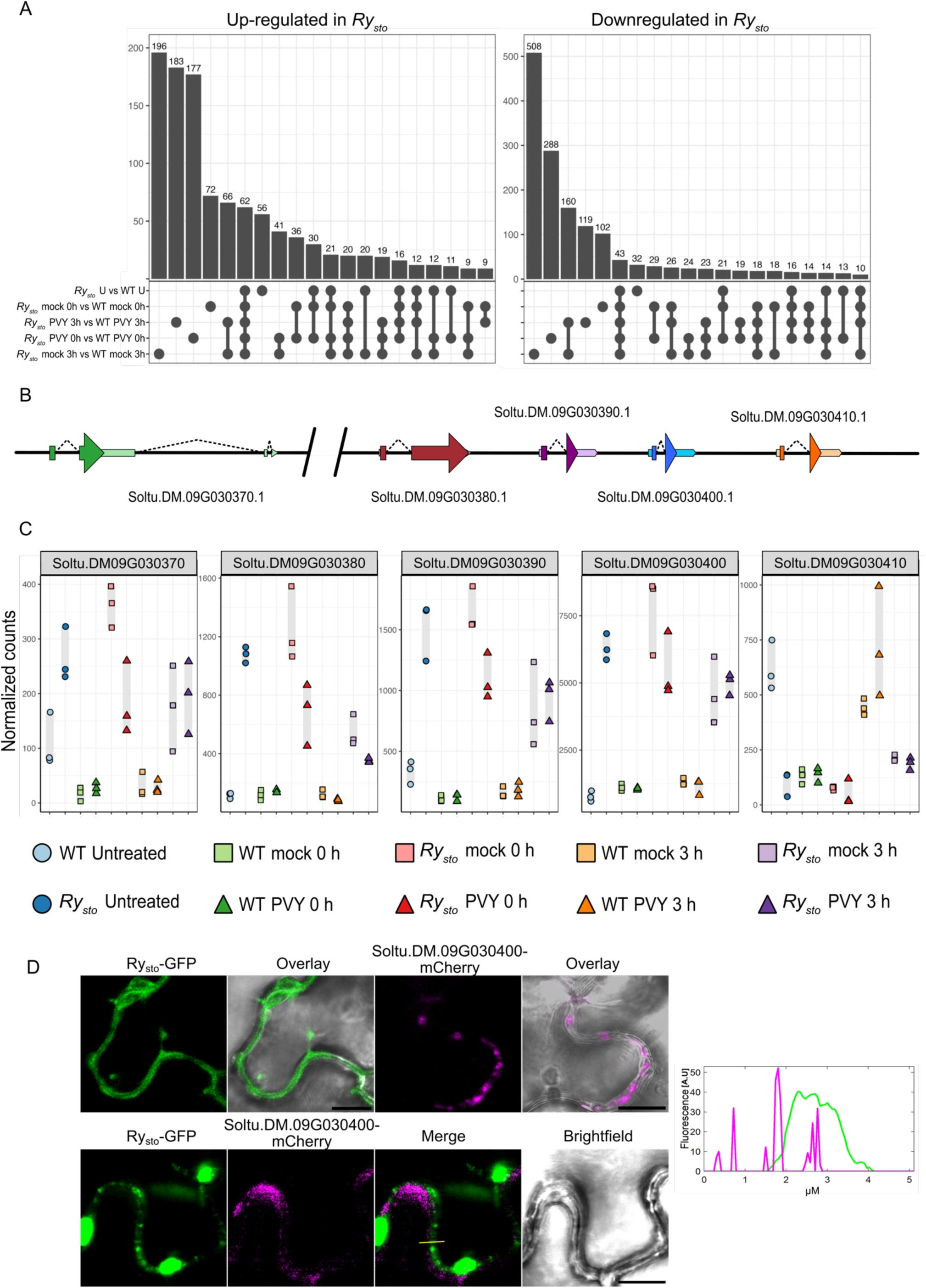
*Ry_Sto_* plants have higher consItuIve expression of genes coding for glycine rich proteins (GRPs) (A) Upset plots depicting the overlaps of up-regulated (lex) and down-regulated (right) genes of all the respective comparisons between *Ry_sto_* and WT Russet Burbank plants. The vertical bars represent the number of genes in each intersection as indicated by the connected black dots. Genes were considered differentially expressed if they met a threshold of |log_2_ fold-change| > 1 and a p-adjusted < 0.05. (B) Gene structures of *Soltu.DM.09G030400* and its orthologs were obtained from *S. tuberosum* genome DM 1-3 516 R44 – v6.1. Orthologs of *Soltu.DM.09G030400* were identified with Orthofinder. The thick colored boxes and arrows represent the coding sequences and thin colored ones represent the UTRs. The dashed lines indicate the introns. The black line represents the 28 kb genomic region located in chromosome 9. (C). Dot-plot of normalized counts of the *GRP* genes showing different expression patterns between *Ry_sto_* transgenic and WT plants. The shapes represent the treatments: untreated (U), mock, or PVY inoculation. The colors show the genotype/wme-point combinations. The counts were normalized with the DEseq2 package in R. (D) Cortical view of *N. benthamiana* leaf epidermal cells transiently expressing Ry_sto_-GFP and Soltu.DM.09G030400-mCherry alone (upper panel) and both Ry_sto_-GFP/ Soltu.DM.09G030400-mcherry (lower panel). The fluorescence intensity along the yellow line in the merged image (lex to right) is plotted. The graph illustrates a partially overlapping subcellular distribution of Ry_sto_-GFP and Soltu.DM.09G030400-mCherry at the interface between the cell wall and cytoplasm. Scale bars: 10 µm

Overall, these results demonstrate that *Ry_sto_* plants exhibit a distinct pattern of constitutively expressed genes. This group comprises a set of genes encoding GRPs, which are likely secreted and function in the extracellular matrix.

We cloned the sequence encoding Soltu.DM.09G030400 with a C-terminal mCherry tag and transiently expressed it in *N. benthamiana* to examine its subcellular localization. The protein was predominantly localized to the cell wall; its deposition was not uniform, leading to local enhancements in cell wall thickness (Fig. 2D). To test a potential functional relationship between Soltu.DM.09G030400 and Ry_sto_, we transiently expressed Ry_sto_-GFP either alone or together with Soltu.DM.09G030400-mCherry. When expressed alone, Ry_sto_-GFP was mainly cytoplasmic (Fig. 2D). Co-expression of Soltu.DM.09G030400–mCherry altered Ry_sto_ localization, leading to the formation of large cytoplasmic clusters and smaller condensates at the plasma membrane–cell wall interface. These structures partially colocalized with Soltu.DM.09G030400–mCherry (Fig. 2D). In regions enriched with Soltu.DM.09G030400, Ry_sto_–GFP formed transverse clusters aligned parallel to each other in the cortical region (Fig. 2D). These observations suggest that, in the endogenous potato context, Ry_sto_ may localize near potential virus entry sites.

### Ry_sto_ moves toward the plasma membrane- cell wall continuum upon perceiving an infection

To determine whether virus presence may affect Ry_sto_’s subcellular localization, we infected *N. benthamiana* plants with the PVY-GFP infectious clone and two other GFP-tagged viruses recognized by Ry_sto_, that is TuMV and PPV. Then, we transiently expressed Ry_sto_-RFP in both infected and healthy control plants. In all tested viral backgrounds, we observed changes in Ry_sto_ localization, with symmetric condensates near the plasma membrane and cell wall-interface. This pattern closely resembled the distribution observed for Ry_sto_ co-expressed with the Soltu.DM.09G030400 glycine-rich protein (Fig. 3A). To check whether indeed a subpool Ry_sto_ is relocalized after activation towards plasma membrane and cell wall-continuum, we compared Ry_sto_-FLAG distribution in PVY- infected and healthy plants. To this end, we performed membrane fractionation and apoplastic fluid collection. In samples derived from infected plants, Ry_sto_-FLAG was clearly enriched in pellet P fraction when compared to healthy plants (Fig. 3B) In parallel experiment, we analyzed apoplastic fluid collected from PVY-infected and healthy plants expressing Ry_sto_–HIS-FLAG or YFP–STREP-FLAG (control). Following His-tag affinity purification, the enriched apoplastic proteins were immunoprecipitated using anti-FLAG beads and analyzed with antibodies. Ry_sto_ was detected only in the apoplastic fractions from PVY-infected samples (Fig. 3C), indicating that upon infection a subpool of Ry_sto_ is mobilized into the apoplast (Fig. 3C).

**Figure 3.**
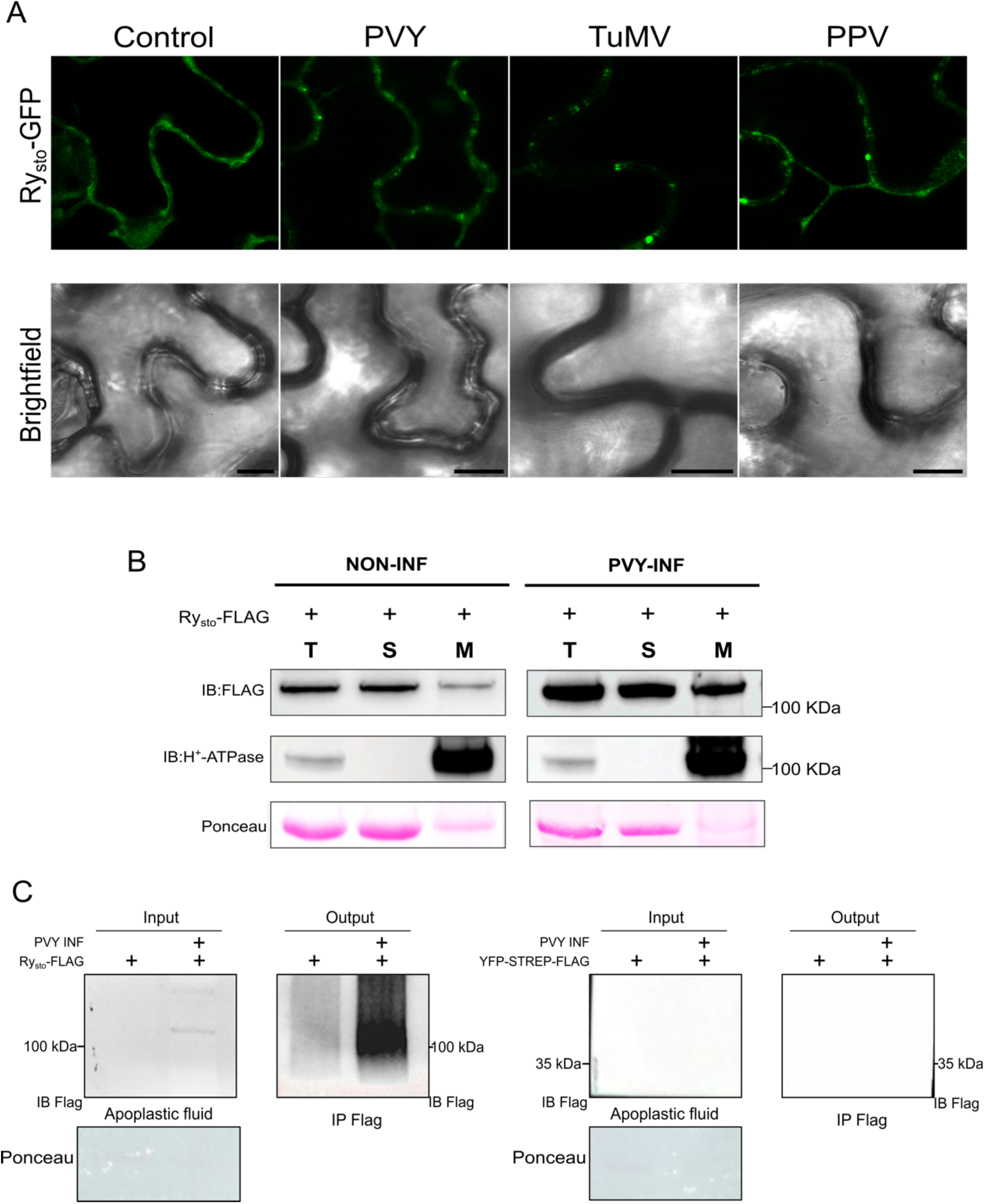
Ry_sto_ partially relocates towards the cell wall and is detectable in the apoplast following viral infection. (A) *Nicotiana benthamiana* plants were agroinfiltrated with infectious GFP-tagged clones of PVY, TuMV, or PPV. Seven days post-infection, when symptoms were visible and GFP fluorescence was detected, systemic leaves were agroinfiltrated with a dexamethasone-inducible Ry_sto_-RFP construct. As a control, Ry_sto_-RFP was infiltrated into healthy, non-infected *N. benthamiana* plants. Twenty hours after induction, confocal microscopy was performed to assess Ry_sto_ subcellular localization. (B) Redistribution of Ry_sto_ subpool upon PVY infection toward the plasma membrane. *N. benthamiana* plants were mechanically inoculated with PVY. Seven days after infection, when viral symptoms were visible, systemic leaves were infiltrated with *Agrobacterium* carrying Ry_sto_-HIS-FLAG. Lysates of harvested samples were fractioned into total (T), soluble (S) and pellet (P) fractions. (C) *N. benthamiana* plants were mechanically inoculated with PVY. Seven days after infection, when viral symptoms were visible, systemic leaves were infiltrated with *Agrobacterium* carrying Ry_sto_-HIS-FLAG or YFP-STREP-FLAG (control). In parallel, the same constructs were infiltrated into healthy, non-infected plants. Forty-eight hours after agroinfiltration, apoplastic sap was extracted using a vacuum infiltration–centrifugation method, and then pooled across samples. Ry_sto_-HIS-FLAG was initially enriched by His-tag affinity purification, and then enriched apoplastic protein fraction was immunoprecipitated using anti-FLAG antibody-conjugated beads. Bound proteins were eluted with 3×FLAG peptide and analyzed by immunoblotting with the appropriate antibodies. Ponceau staining served as a loading control. Chloroplast contamination was quantified by extracting the samples with 80% ethanol, followed by spectrophotometric analysis at 600 nm. Readings at A_600_ for apoplastic fluid were undetectable.

### Host Hsp70 and CPIP cochaperonins are required for Ry_sto_ activation

The constitutive enrichment in cell wall–associated and defense-related genes observed in *Ry_sto_* plants, together with the early and extensive transcriptional reprogramming, suggests that ER depends on rapid coupling of virus perception with intracellular signaling. We hypothesized, that a key candidate for mediating this process is the host chaperone machinery, which plays a central role in PVY entry, intracellular trafficking, and cell-to-cell movement (Hoffius et la, 2007; Alazem et al., 2025). Therefore, we checked the expression pattern of *St*Hsp70 (*Soltu. DM.06G031410*) containing a conserved motif required for plasmodesmatal trafficking (Aoki et al., 2002), and *St*CPIP isoforms (*Soltu.DM.04G001130* and *St*CPIP2a *Soltu. DM.05G006580*). Interestingly, both Hsp70 and CPIP1 were upregulated at 3 h in *Ry_sto_* and WT plants, with the upregulation being stronger in *Ry_sto_* plants (Fig 4A). In contrast, CPIP2a was downregulated. Although these changes were not high, given the localized character of ER they were significant (Figure 4B).

**Figure 4.**
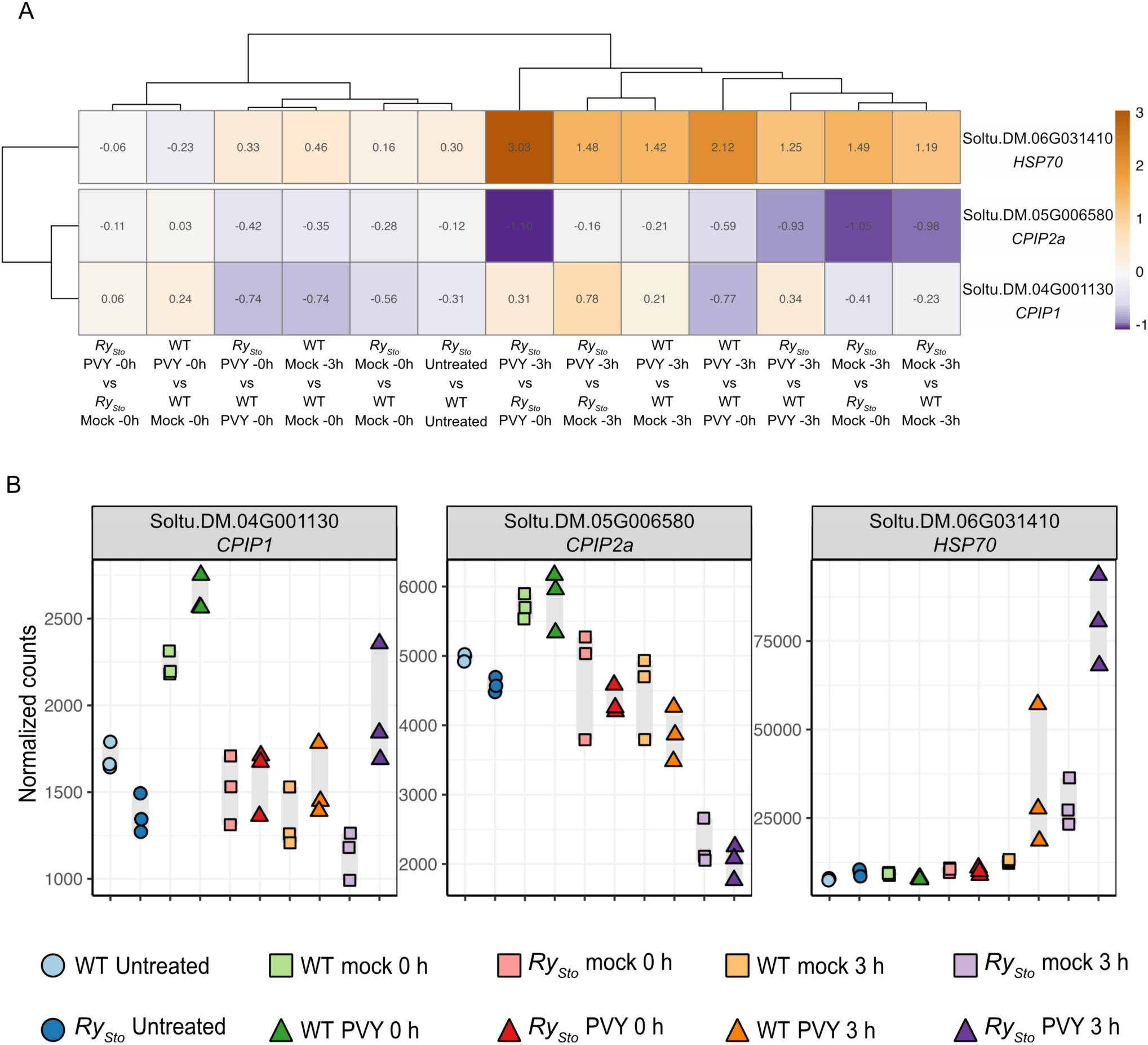
DifferenIal expression of *Hsp70*, *CPIP1*, and *CPIP2a* in *Ry_sto_* and wild-type *Russet Burbank* plants. (A) Hierarchical heatmap showing the relative expression of *StHsp70*, *StCPIP1* and *StCPIP2a* calculated with the DESeq2 package. The lateral bar indicates the log2 fold change. The *S. tuberosum* genome DM 1-3 516 R44v6.1 was used to obtain ortholog sequences. (B) Dot plot of normalized counts for *StHsp70*, *StCPIP1,* and *StCPIP2a* genes, revealing different expression patterns between *Ry_sto_* and WT plants. The shapes represent treatments: untreated (U), mock, or PVY inoculated. Colors indicate genotype/wme-point combinations. Counts were normalized using the DESeq2 package in R.

We therefore hypothesized that Ry_sto_ detection system may rely on the viral dependency of the same Hsp70–CPIP network during early infection steps, thus converting a susceptibility pathway into a signal for defense activation.

We cloned the corresponding *StHsp70*, *StCPIP1*, and *StCPIP2a* and tested their roles in Ry_sto_ function. Of not, we previously found that within *N. bethamiana* system, Ry_sto_ shifts the response from ER to HR (Grech-Baran et al., 2022).

Silencing of endogenous CPIP1 and CPIP2a in *N. benthamiana* by VIGS (Kumar and Mysore, 2014). significantly reduced Ry_sto_-triggered hypersensitive response (HR) upon PVY CP co-expression (Fig. 5A–C). Consistently, complementation with potato-derived CPIP1 or CPIP2a restored HR, with the strongest effect seen for CPIP1, as assessed by HR scoring and ion leakage tests (Fig. 5A, C). In contrast, silencing did not affect HR initiated by the unrelated TNL Roq1 upon detection of the cognate effector XopQ (Supplemental Fig. 1) indicating the necessity of chaperonins specificallly for Ry_sto_ signaling. As expected, in the silencing control plants, that is infiltrated with TRV2-GFP, HR establisment mediated by Ry_sto_ and CP was not compromised (Fig 5A).

**Figure 5.**
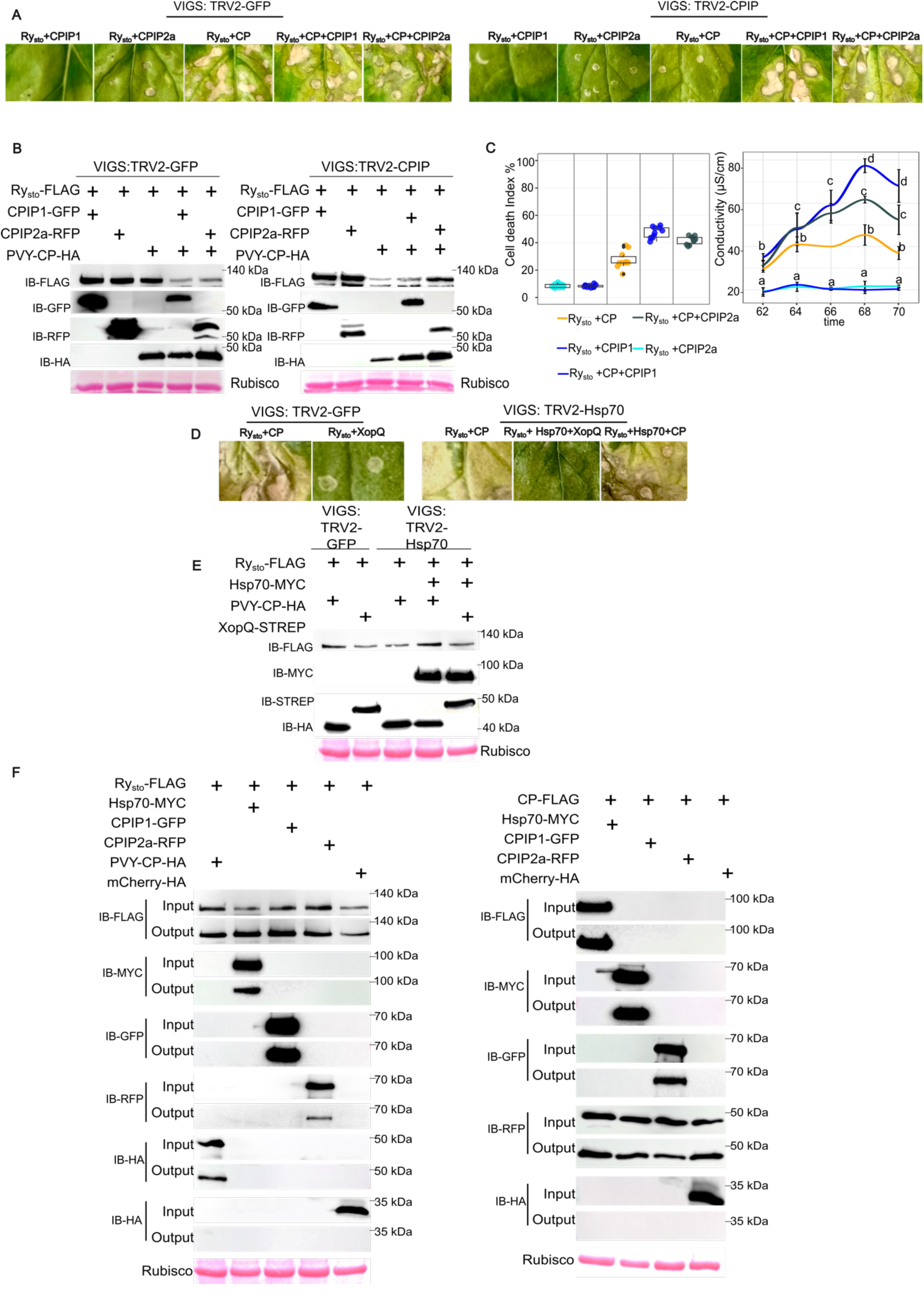
Hsp70 and CP-interacting cochaperonins are essential for Rysto-mediated immune activation. (A–C) Silencing of CPIP cochaperonins impairs Ry_sto_-mediated hypersensitive response. Leaves of *N. benthamiana* plants, silenced through VIGS with TRV2-GFP (control, left) or TRV2-CPIP (right), were co-infiltrated with constructs encoding Ry_sto_-FLAG, PVY coat protein (CP-HA), and the potato-derived cochaperonins *St*CPIP1-GFP and/or *St*CPIP2a-RFP in the specified combinations. (A) Representative hypersensitive response (HR) phenotypes. (B) Immunoblot analysis showing the accumulation of expressed proteins. (C) HR scoring (left) and ion leakage measurement (right). (D–E) Hsp70 is required for Ry_sto_-dependent HR induction. TRV2-GFP (control)– or TRV2-Hsp70–silenced *N. benthamiana* leaves were co-infiltrated with Ry_sto_-FLAG and PVY CP-HA, solely or complemented with potato-derived *St*Hsp70-MYC. XopQ-STREP was included as an unrecognised control. (D) Representative HR phenotypes. (E) Immunoblot analysis confirming protein accumulation. (F) Both Ry_sto_ and PVY CP interact with *St*Hsp70 and cochaperonins network *in planta*. Leaves of *N. benthamiana* plants were co-infiltrated with CP-FLAG together with *St*Hsp70-MYC, *St*CPIP1-GFP, *St*CPIP2a-RFP, or mCherry-HA, or with Ry_sto_-FLAG along with CP-HA, *St*Hsp70-MYC, *St*CPIP1-GFP, *St*CPIP2a-RFP, or mCherry-HA. Protein complexes were immunoprecipitated using anti-FLAG–conjugated beads, eluted with 3×FLAG peptide, and analyzed by immunoblotting with the indicated antibodies. Ponceau staining is shown as a loading control. Experiments were performed with at least three biological replicates. Protein extraction and immunodetection were conducted at 3 days post-infiltration.

Similarly, silencing of endogenous Hsp70 suppressed Ry_sto_-mediated HR, and complementation with *St*Hsp70 restored lesion formation (Fig. 5D, 4E). Control experiments were conducted again in TRV2-GFP–“silenced” plants and using Roq1 – XopQ system (Fig. 5, Suppl. Fig. 2). Immunoblotting confirmed appropriate protein expression.

Collectively, Hsp70 and CPIP cochaperonins are essential for efficient Ry_sto_ activation, supporting a model in which components of the viral movement machinery contribute to immune activation.

### Ry_sto_ interacts with Hsp70 and CPIPs and forms a pre-activation complex

Having established that Ry_sto_ physically interacts with PVY CP (Grech-Baran et al., 2022), we now investigated whether Ry_sto_ engages the same host chaperone network as PVY CP does. To this end, we performed in planta co-immunoprecipitation assay (Fig. 5F). PVY CP-FLAG was found to associate with *St*Hsp70-MYC, *St*CPIP1-GFP, and *St*CPIP2a-RFP, consistent with previous reports (Hoffius et al., 2007) (Fig. 5F). Ry_sto_-FLAG was co-expressed along with each of these candidate partners, with CP-HA and mCherry serving as positive and negative controls, respectively. Co-immunoprecipitation followed by immunoblot analysis revealed that Ry_sto_-FLAG co-purified with *St*Hsp70 and both *St*CPIP isoforms (Fig. 4F), indicating that Ry_sto_ participates in this host factor network. Importantly, PVY CP demonstrated a significantly stronger association with Hsp70 than Ry_sto_, while both proteins showed a higher affinity for CPIP1 than for CPIP2a (Fig. 5F).

To investigate the organization of the Ry_sto_–Hsp70–PVY-CP–CPIP network, we performed Blue Native PAGE (BNP), which detects native protein complexes (Eubel et al., 2005). To this end, Ry-FLAG was transiently co-expressed with *St*Hsp70-MYC, or with *St*Hsp70-MYC and PVY-CP-HA in *eds-1* knockout plants. As controls, Ry_sto_-FLAG was co-expressed with PVY-CP alone or with mCherry.

Co-expression of Ry_sto_ with *St*Hsp70 resulted in the detection of a native complex of approximately 480 kDa, corresponding to a pre-active Ry_sto_–Hsp70 complex (Fig. 6A, B). Whereas co-expression with PVY-CP triggered the formation of a high-molecular-weight complex exceeding 720 kDa, corresponding to Ry_sto_ oligomerization and assembly of active resistosomes (Fig. 6A).

**Figure 6.**
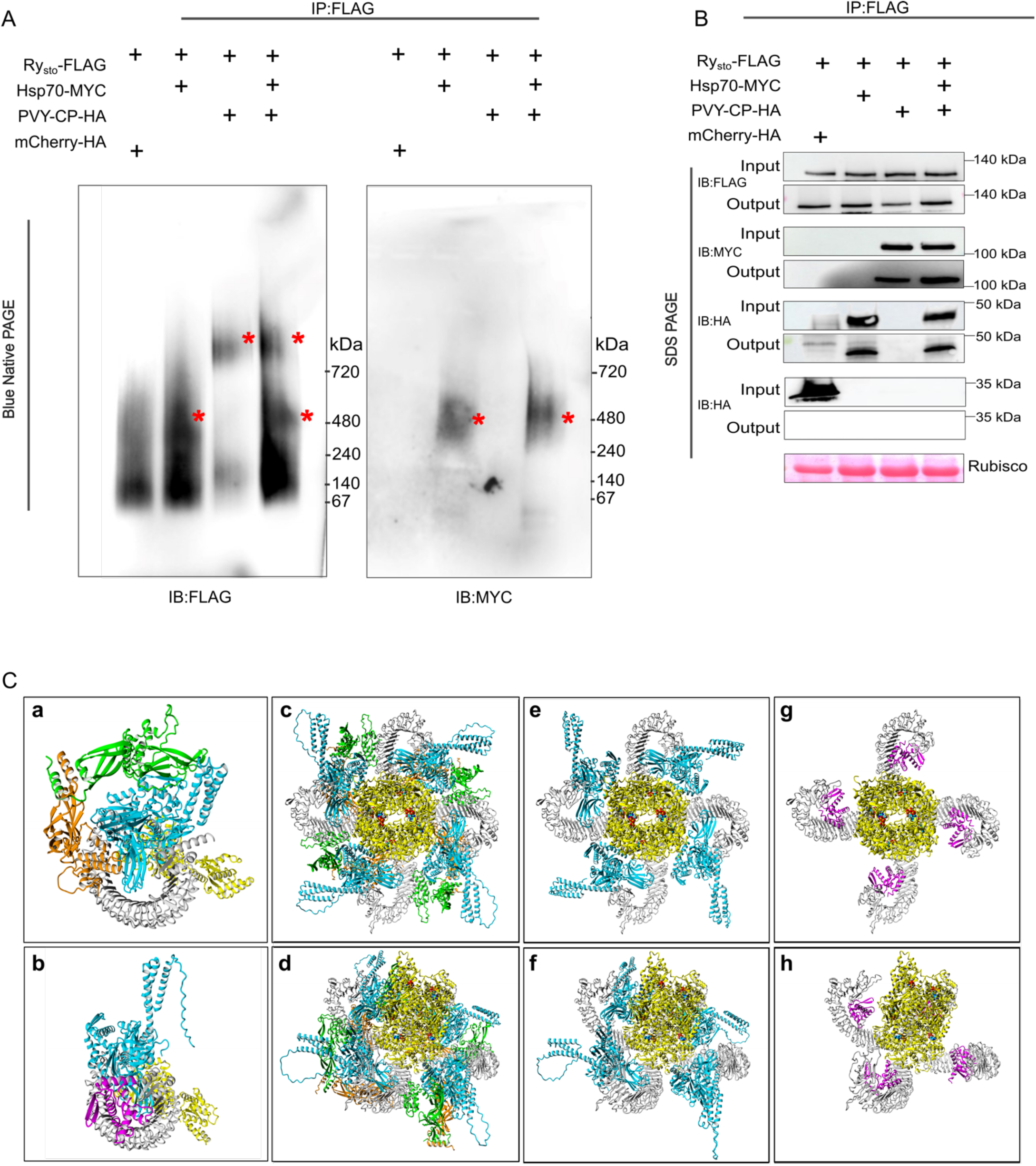
Ry_sto_ forms a pre-active complex with *St*Hsp70 and oligomerizes into an active resistosome upon PVY CP recognition. (A) Pre-active dimers and CP-dependent oligomerization of Ry_sto_. Leaves of *Nicotiana benthamiana* were co-infiltrated with Ry_sto_-FLAG, *St*Hsp70-MYC, CP PVY-HA, and mCherry-HA as a negative control. Two days post-infiltration, total protein extracts were prepared, and samples were immunoprecipitated using anti-FLAG–beads. Eluted proteins were subjected to blue native PAGE (BNP) and immunoblotted with anti-FLAG and anti-MYC antibodies. Ry_sto_ co-migration with Hsp70 at ∼480 kDa corresponds to a pre-active Ry_sto_–Hsp70 complex, whereas higher-molecular-weight complexes over 720 kDa detected in the presence of PVY CP indicate resistosome formation. (B). The same samples used for BNP were analyzed by SDS-PAGE followed by immunodetection with the appropriate antibodies to confirm the presence of Ry_sto_-FLAG and co-precipitated partners. Ponceau S staining is shown as a loading control. Experiments were performed with at least three biological replicates. (C) Structural model of Ry_sto_ activation by PVY coat protein. The LRR domain of Ry_sto_ is shown in grey, the TIR and NB domains in yellow, PVY CP in magenta, Hsp70 in blue, and the co-chaperones CPIP1 and CPIP2a in orange and green, respectively. Structural models of Ry_sto_ in complex with Hsp70 and its co-chaperones (a) or with PVY CP (b) support a competitive mode of ligand binding, as both ligands are predicted to interact with an overlapping surface on the LRR domain, making their binding mutually exclusive. Structural modeling further indicates that the rigid domain of Hsp70 is considerably larger than PVY CP, rendering its accommodation within the homology-modeled tetrameric active conformation of Ry_sto_ unfavorable (c,d). Additional binding of CPIP1 and CPIP2a leads to significant steric clashes, further destabilizing the tetrameric receptor assembly (e,f). In contrast, PVY CP binding is compatible with the tetrameric form of Ry_sto_ and does not introduce steric hindrance, thereby facilitating receptor activation (g,h).

Interestingly, when Ry_sto_ was co-expressed with both PVY-CP and *St*Hsp70, the active oligomeric complex was detected at a similar molecular weight; however, a distinct fraction of the ∼480 kDa complex remained visible upon FLAG immunodetection. Notably, MYC antibodies failed to detect the high-molecular-weight oligomer, indicating that *St*Hsp70 is not stably associated with the active resistosome (Fig. 6A). These data suggest that the Ry_sto_–Hsp70 interaction occurs predominantly before Ry_sto_ oligomerization. Although we cannot exclude that lack of *St*Hsp70 detection is due to structural restrictions.

Consistent with this interpretation, structural homology modeling and AI-assisted predictions (Krieger et al., 2009), along with Co-IP data (Fig. 5F), suggest that PVY CP binds to Ry_sto_ by competing with Hsp70 and CPIPs. Both ligands target a similar region of the LRR domain, preventing simultaneous binding (Fig. 6C). While the binding of Hsp70 is stabilized by CPIP1 and CPIP2a (Fig. 6C), the large, rigid Hsp70 molecule cannot fit into the homology-modeled tetrameric active form of Ry_sto_, and additional CPIP binding causes significant steric clashes (Fig. 6C). In contrast, PVY CP binding does not cause steric interference and is compatible with tetrameric receptor activation. We propose that PVY CP binding to the receptor–chaperone complex displaces Hsp70, switches Ry_sto_ from its inactive form, and facilitates resistosome oligomerization and activation.

## Discussion

Enhancing endogenous antiviral defenses is essential for reducing crop losses, and extreme resistance offers a durable strategy for virus control (Sett et al., 2022). Our study shows that Ry_sto_-mediated ER involves rapid PVY perception, early transcriptional reprogramming, and coordinated activation of metabolic and redox-associated defenses. Together with the observation that viral infection promotes relocalization of the receptor toward the plasma membrane-cell wall interface, and biochemical evidence that the host chaperone network used by PVY virus is required for Ry_sto_ function, these findings support a model in which early recognition events are quickly translated into systemic responses that restrict viral spread.

Transcriptome analysis revealed extensive host gene reprogramming in *Ry_sto_* plants as early as 3 h post-inoculation (Fig. 1). Of note, PCA analysis separated Ry_sto_-expressing and WT Russet Burbank plants already before infection, indicating that a transcriptional priming contributes to ER phenomenon (Fig. 2). At 3 h post-inoculation, *Ry_sto_* plants exhibited nearly three-fold more differentially expressed genes than WT plants, highlighting the importance of short response time and high amplitude for an effective antiviral immunity (Fig. 1). Functional enrichment analysis showed that *Ry_sto_*-specific up-regulated genes are associated with glutathione metabolism and aromatic amino acid biosynthesis, linked to redox balance, ROS signaling, and the production of defense-related secondary metabolites, including phenylpropanoids, salicylic acid and pathogenesis-related protein precursors (Fig. 1). An activation of a similar set of defense mechanisms was observed in *Ny-1*–mediated PVY resistance that is accompanied with a classic HR, however, at the later infection stages (Beabler et al., 2014). Although ROS involvement in ER has not been previously described, ROS-induced symptomless TMV resistance in tobacco has been reported (Kuenstler et al., 2016). In contrast, susceptible wild-type plants displayed a limited early transcriptional response, with low level activation of defense-related pathways, reflecting delayed or insufficient reprogramming that allowed PVY replication and spread (Fig. 1).

*Ry_sto_* plants also exhibited a constitutive defense-related transcriptomic profile, including elevated expression of 12 pathogenesis-related genes, WRKY and TGA transcription factors, and other immune-associated genes (Fig. 2). TGA, particularly the Golden 2-like transcription factor involved in ABA signaling, has been implicated in Rx1-mediated ER against PVX (Ahmad et al., 2019). Notably, a cluster encoding glycine-rich proteins (GRPs) on chromosome 9, including Soltu.DM.09G030400, was highly expressed in *Ry_sto_* plants. Transient expression in *N. benthamiana* confirmed cell wall localization and induced structural remodeling, and co-expression with Ry_sto_ led to receptor aggregation near the plasma membrane and cell wall (Fig. 2). These findings suggest that Ry_sto_ may be pre-positioned at the cell periphery—the first barrier to viral invasion—and that secreted GRPs may facilitate early immune positioning and rapid PVY recognition. The positive role of GRPs was also described for Rx, where NbGRP7 associates *in planta* with this receptor and positively regulates Rx1-mediated extreme resistance (Sakura et al., 2022). In *Nicotiana glutinosa*, the expression of a gene encoding a glycine-rich RNA-binding protein (*Ng*RBP) was negatively regulated by TMV infection (Naqvi et al., 1998). A role of priming/activating the innate immune response was described for *Arabidopsis thaliana* Pep1, which, by activation of transcription of the defensive gene (*PDF1.2*), activates the synthesis of H_2_O_2_ (Huffaker et al., 2006).

The enrichment of the transciripts encoding secreted GRPs, including *Soltu.DM.09G030400*, and the ability of its product to reposition Ry_sto_ toward the cell wall suggests that transcriptional priming creates a cellular environment in which Ry_sto_ is spatially aligned with Hsp70-dependent viral movement complexes. Consistent with this model, in response to PVY, TuMV, and PPV, Ry_sto_ accumulated in the condensates at the inferface of the plasma membrane and cell wall. This suggests that the redistribution of Ry_sto_ upon infection is a general step of Ry_sto_ activation (Fig. 3). The similarity between virus-induced localization of the receptor and the pattern observed upon its co-expression with the glycine-rich protein Soltu.DM.09G030400 supports a functional link between extracellular components and Ry_sto_ positioning. Importantly, the detection of Ry_sto_ in the apoplast specifically after PVY infection indicates that a subpool of Ry_sto_ can be mobilized beyond the cytoplasm (Fig. 3). Such relocalization of an intracellular immune receptor to the cell periphery may enhance early pathogen sensing or signal initiation at sites of viral entry and cell-to-cell movement, thereby contributing to the rapid and effective ER execution. Fascinatingly, a similar pattern of localization and virus perception was observed for Rsc4-3, which encodes a cell-wall-localized NLR-type resistance protein conferring resistance to soybean mosaic virus (Yin et al., 2021).

Ry_sto_-mediated ER response relies, at the molecular level, on a specific network of Hsp70- and CP-interacting cochaperonins (CPIPs) that prime the receptor for rapid viral detection (Fig. 4). Silencing for Hsp70 or CPIPs in *N. benthamiana* strongly suppressed Ry_sto_-triggered immune response, whereas complementation with potato-derived Hsp70 or CPIPs restored immune activation, confirming the specificity and crucial role of this module (Fig. 5). The same Hsp70-cochaperonin network is used by PVY during cell entry (Hoffius et al., 2007; Hafrén et al., 2010). The interaction with Hsp70 takes place in plasmodesmata to facilitate the transport of movement-competent viral ribonucleoprotein complexes (Aoki et al., 2002; Hoffius et al., 2007). The interaction of Ry_sto_ with Hsp70, along with its rapid partial re-localization toward the plasma membrane and cell wall upon infection, suggests that Ry_sto_ activation occurs near plasmodesmata, which can serve as immune signaling checkpoints. This spatial repositioning may enable efficient sensing of PVY coat protein–Hsp70 complexes and contribute to the rapid restriction of viral spread characteristic of extreme resistance. Thus, Hsp70 plays a dual role as both a susceptibility factor and a crucial regulator of immune activation during Ry_sto_-PVY interaction. A similar role of Hsp70 was recently described in *A. thaliana* during infection with *Plasmodiophora brassicae* (<u>Kopecká</u> et al., 2025).

Further analysis showed that Ry_sto_ forms a pre-active complex with Hsp70, CPIP1, and CPIP2a, stabilizing the receptor in an inactive dimeric state (Fig. 5, 6). Blue Native PAGE demonstrated that PVY coat protein (CP) induces Ry_sto_ oligomerization into a high-molecular-weight resistosome, whereas Hsp70 was not detected in the activated complex, indicating its primary function in the pre-active conformation (Fig. 6). The absence of Hsp70 in the activated complex indicates that its primary function is receptor stabilization prior to activation, consistent with the studies on other plant NLR resistosomes (Wang et al., 2019; Martin et al., 2020). The fact, that in the co-immunoprecipipation assay PVY CP bound more efficiently to HSP70 than Ry_sto_ did suggested that both proteins differ significantly in their binding affinity for HSP70. This allowed us to formulate a hypothesis, that CP outcompetes Ry_sto_ for HSP70, which is in line with the structural modelling data (Fig. 6). We therefore propose that PVY-CP binding to Hsp70 in the receptor complex, detaches the chaperone from Ry_sto_ enabling the transition from the pre-active conformation to the active resistosome (Fig. 6).

In summary, our findings reveal a chaperone-driven mechanism that prepares Ry_sto_ for quick and targeted antiviral defense. By integrating immune priming, spatial receptor repositioning, and chaperone-dependent control, Ry_sto_ showcases how plants can exploit viral host dependencies to achieve fast and durable resistance.

## Materials and methods

### Plant material

*N. benthamiana* plants were grown for 6 weeks in soil under controlled environmental conditions (16 h light / 8 h dark) under permissive (22°C) temperature as described previously (Hoser et al., 2013). Transgenic *Russet Burbank and WT* plants were obtained as described (Grech-Baran et al., 2020). Plants were grown for 4 weeks in soil under controlled environmental conditions (22°C, 16-h light; 18°C, 8-h dark) as described previously (Szajko et al., 2008).

### Constructs assembly

*Ry_sto_* and *Roq1*, *PVY-CP*, *s.t. Hsp70*, *s.t. CPIP1*, *s.t CPIP2a* and *Soltu.DM.09G030400* constructs were assembled using the Golden Gate Modular Cloning (MoClo) system (Weber et al., 2011), along with the MoClo Plant Parts Kit (Engler et al., 2014). The dexamethasone-inducible RY-RFP and mCherry-HA were cloned into the entry pENTR/D-TOPO vector. The resulting entry clones were LR recombined with the Gateway binary pBAV vectors. The constructs are listed in the Suppl. Table 3.

### Transient expression assay

Binary constructs for transient expression were transformed into *Agrobacterium tumefaciens* strain GV3101 (pMP90). *Agrobacterium* cells were resuspended in an infiltration buffer (10 mM MgCl₂ and 10 mM MES, pH 5.6, 100 μM acetosyringone). The optical density at 600 nm (OD₆₀₀) of each construct-containing *Agrobacterium* suspension was adjusted to 0.5, except for *Agrobacterium* containing p19, where it was adjusted to 0.2. Infiltration suspensions were incubated for 1-2 hours at room temperature prior to infiltration. Hypersensitive response (HR)-related phenotypes were evaluated at three days post-infiltration (dpi).

### HR scoring

For the HR assay, 4-week-old *N. benthamiana* plants were used. Constructs in *Agrobacterium* were infiltrated or co-infiltrated into the abaxial surface of *N. benthamiana* leaves. The HR phenotypes were scored at 5 dpi. Detached leaves were photographed under UV light with a yellow filter (Wratten K2 Yellow Filter no. 8; Kodak) attached to the camera lens. Images were captured with a 2-second-long exposure, F8.0, and ISO 400, then scored using ImageJ software according to the scale described in Grech-Baran et al., 2022.

### Ion conductivity

At the specified time points, eight leaf discs (1 cm in diameter) were cut from infiltrated zones and floated abaxial side up on 5 ml of MilliQ water for 10 minutes at 18°C with gentle stirring (50 rpm). The conductivity of the water was measured using a WTW InoLab Multi 9310 IDSCDM83 benchtop meter and reported in μS cm^−1^.

### mRNA preparation and sequencing

To determine which genes are differentially expressed in transgenic *Solanum tuberosum* cv. For Russet Burbank plants expressing the *Ry_sto_* gene, we performed comparative transcriptomic profiling at three time points during early infection. For each factor combination, we prepared three biological repeats. Total RNA from leaf tissue was extracted from *S. tuberosum* cv. Russet Burbank using the RNeasy Plant Mini Kit followed by RNA Clean & Concentrator (Zymo Research). RNA quality assessment, library preparation, and sequencing were performed by Novogene Co., Ltd. (Cambridge, UK). For library preparation, mRNA was enriched from total RNA using poly-A selection. Sequencing was performed on an Illumina NovaSeq X Plus platform, generating approximately 40 million 150 bp paired-end (PE150) reads per sample.

### Protein extraction

For protein extraction, samples were collected from *Agrobacterium*-infiltrated leaves at two dpi using a cork borer. All leaf discs were transferred into 2 ml Eppendorf tubes and rapidly frozen in liquid nitrogen. The samples were then ground into a powder. A protein extraction buffer containing 50 mM Tris-Cl (pH 7.5), 150 mM NaCl, 10% glycerol, 1 mM EDTA pH 8.0, 5 mM DTT, 0.2% IGEPAL, and protease inhibitor cocktail was vortexed with the ground tissue. Centrifugation at 18,000 g for 15 minutes, followed by 5 minutes, was used to remove cell debris. Aliquots of the samples were either used directly for Blue Native-PAGE (BNP) or subjected to immunoprecipitation (IP) on Flag beads, followed by BNP or SDS-PAGE. For SDS-PAGE, aliquots of the input and Co-IP samples were mixed with 3X SDS sample buffer (containing 30% glycerol, 3% SDS, 93.75 mM Tris-Cl pH 6.8, and 0.06% bromophenol blue) with 100 mM DTT and heated at 70°C for 10 minutes.

### BNP and SDS-PAGE analysis

For blue native polyacrylamide gel electrophoresis (BNP), clarified protein extracts or IP eluates were diluted according to the manufacturer’s instructions with NativePAGE™ 5% G-250 sample additive, 4× NativePAGE Sample Buffer. Samples were loaded onto 3%–12% Bis-Tris NativePAGE™ gels (Invitrogen) and ran alongside SERVA Native Marker (SERVA). Following the separation, proteins were transferred to polyvinylidene difluoride (PVDF) membranes using NuPAGE™ Transfer Buffer and the Trans-Blot Turbo Transfer System (Bio-Rad), according to the manufacturer’s protocol. Membranes were fixed with 8% acetic acid for 15 minutes, rinsed with water, and air-dried. To visualize native protein standards, membranes were reactivated with ethanol prior to immunoblotting. For SDS-PAGE, protein samples were mixed with SDS loading buffer and denatured at 72°C for 10 minutes. Proteins were transferred to PVDF membranes using the Trans-Blot Turbo Transfer System with the supplied transfer buffer, following standard procedures.

### Immunoblotting and detection

Membranes were blocked in 5% (w/v) non-fat dry milk in Tris-buffered saline containing 0.01% Tween-20 (TBS-T) for 1 hour at room temperature. Blots were then incubated overnight at 4°C with horseradish peroxidase (HRP)-conjugated primary antibodies diluted 1:5,000 in 5% milk / TBS-T. The following antibodies were used: anti-GFP (B-2, Santa Cruz Biotechnology), anti-Strep (71591-M, Sigma-Aldrich), anti-Flag (M2, Sigma-Aldrich), anti-RFP (ab183628, Abcam), anti-HA (12013819001-HRP conjugated, Sigma-Aldrich), and anti-Myc (conjugated HRP, Merck clone 9E10). Signal detection was performed using Pierce™ ECL Western Blotting Substrate (32106, Thermo Fisher Scientific), with up to 50% SuperSignal™ West Femto Maximum Sensitivity Substrate (34095, Thermo Fisher Scientific) added for enhanced sensitivity when needed. Chemiluminescence was imaged using Azure 400 AZI400-01 (Azure Biosystems, USA). Membrane loading controls were visualized by staining with Ponceau S (Sigma-Aldrich).

### Confocal laser scanning microscopy

Cortical view of *N. benthamiana* leaf epidermal cells transiently expressing Ry_sto_-GFP and Soltu.DM.09G030400-mCherry alone (upper panel) and both Ry_sto_-GFP/ Soltu.DM.09G030400-mCherry (lower panel). The fluorescence intensity along the yellow line in the merged image (left to right) is plotted. The graph illustrates a partially overlapping subcellular distribution of Ry_sto_-GFP and Soltu.DM.09G030400-mCherry at the interface between the cell wall and cytoplasm. Scale bars: 10 µm.

### Viral Infectious Assays

PVY^-NTN^ isolate (AJ585342.1) infection was performed as described previously (Grech-Baran *et al*., <u>2020</u>). For PVY-GFP, PPV-GFP, and TuMV-GFP infection, plants were infected using PVY-GFP (own disposal), PPV-GFP, or TuMV-GFP infectious clones (Fernández-Fernández *et al*., <u>2001</u>; Garcia-Ruiz *et al*., <u>2015</u>), as described previously (Salvador *et al*., <u>2008</u>; Garcia-Ruiz *et al*., <u>2015</u>).

### Membrane fractionation

*N. benthamiana* plants were mechanically inoculated with PVY. Seven days post-infection, leaves were infiltrated with *Agrobacterium* carrying Ry_sto_-FLAG. Non-infected plants infiltered Ry_sto_-FLAG served as controls. Samples were collected 48 hours post-infiltration. Membrane fractionation was performed as described by Contreras et al. (2022). Proteins were then analyzed by immunoblotting using anti-FLAG antibodies, with anti-H⁺-ATPase as a membrane marker.

### Apoplastic fluid collection

*N. benthamiana plants* were mechanically inoculated with PVY. At seven days post-infection, leaves were infiltrated with *Agrobacterium* carrying either Ry_sto_-His-FLAG or YFP-STREP-FLAG (control). In parallel, non-infected plants were infiltrated with the same constructs. Apoplastic fluid was collected as described by Ishihama et al. (2025), followed by His-tag affinity purification and immunoprecipitation using anti-FLAG–conjugated beads.

### Statistical analysis

Statistical analyses were performed using R 4.5.0 within R Studio 2025.05.0+496. Technical replicates included multiple readings from the same plant in a single experiment, while biological replicates involved measurements from independent plants. Data analysis pipeline was as follows: first, data were tested for suitability for parametric analysis by assessing the normality of residuals with the Shapiro–Wilk test. If the data met the assumptions for parametric testing, a repeated-measures analysis of variance (ANOVA) was conducted, followed by Tukey’s honestly significant difference (HSD) test (**, P < 0.001).

### Structural modelling

Structures of the receptors were initially modeled by homology with the local instance of Yasara Structure package (v. 21.8.27). Up to 8 template structures were analyzed; for each, up to 5 alignment variations were tested. For each modeled loop, up to 50 conformations were tested. The maximal oligomerization state was limited to the tetramer. The quality of the resulting models was assessed with the Z-score value, which distribution along the sequence was monitored to identify the questionable regions of the protein. In most cases, a model built on the EM structure of a disease resistance protein (i.e., 7CRC, 7DFV, 7CRB, 7JLV, or 7JLU) was scored as the best, with a preference for the tetrameric structures build on a shorter template (7DFV), which still covered the regions of interest (WHD and LRR domains). Finally, the best parts of the single-template models were combined into a hybrid model.

The active form of any tested receptor was always assigned to the highest-scoring tetramer. Screening of receptor structures was performed using both the local AlphaFold v2.3.2 instance package and AlphaFold3. We used either Monomer or Multimer versions to model monomers or tetramers.

### Bioinformatic analyses

The quality of the reads was assessed with FastQC. The reads were mapped to the *S. tuberosum* genome DM_1-3_516_R44 v6.1 (Pham *et al*., 2020) using HISAT2 version 2.2.1 (Kim *et al*., 2015). The quantification to retrieve the counts matrix was done with featureCounts version 2.0.6 (Liao *et al*., 2014). The differential expression analysis was performed with DEseq2 version 1.42.0 using a |log2(FoldChange)| >= 1 and p-adjusted value <= 0.05 (Love *et al*., 2014). The mapping of the reads to the PVY genome (NCBI accession number GCF_000862905.1) was performed using Bowtie2 aligner (Langmead & Salzberg, 2012). The prediction of secretion signal and identification of protein motif or domains was done with SignalP and InterProScan respectively (Nielsen *et al*., 2024; Blum *et al*., 2025).

## Acknowlegments

We would like to thank Prof. Juan Antonio Garcia Alvarez and Dr. Véronique Decroocq for sharing PPV-GFP infectious clones. We also thank Prof. James Carrington for sharing the TuMV-GFP infectious clone. The research was supported by a SONATA 15 (2019/35/D/NZ9/03565) grant from the National Science Centre, Poland, awarded to M. G-B.

## Author Contributions

Conceptualization: M.G.-B., J.C. O. Methodology: M.G.-B., J. C. O., J.T.P., T.V.C, M.L., I.B.F. Formal analysis: M.G.-B., M.K., J.C.O. Investigation: M.G.-B., J. C. O., J.T.P., T.V.C, M.L., I.B.F. Writing – Original Draft: M.G.-B., M.K., J.C.O. Writing – Review & Editing: all authors. Visualization: M.G.-B., J.C.O., T. V.C., M.L. Supervision: M.G.-B., M.K.

## Data Availability

The data that support the findings presented in this publication are available from the corresponding author upon reasonable request.

---

**Supplementary Figure 1.**
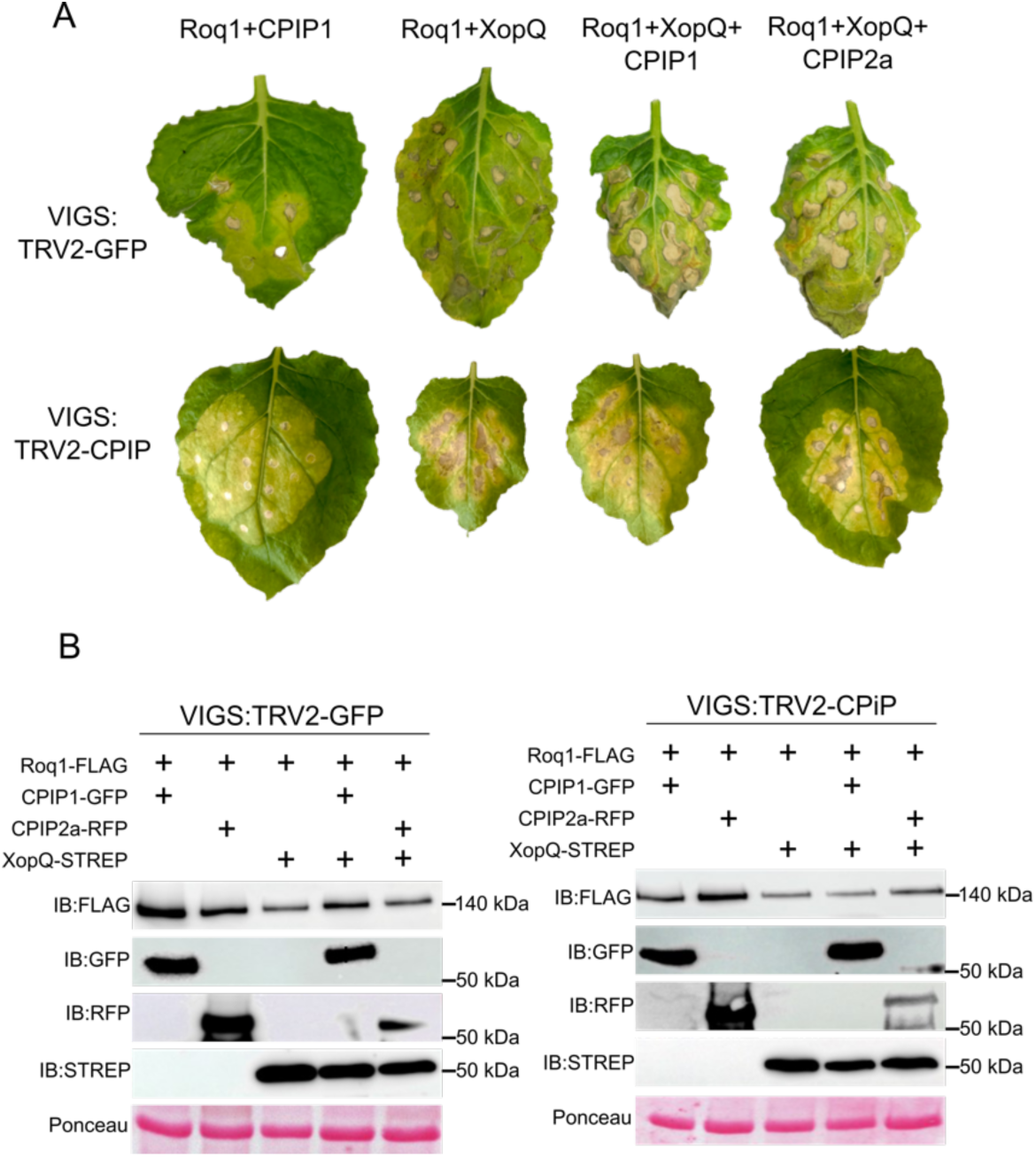
Control analysis of the Roq1–XopQ system in plants silenced for TRV2-GFP or TRV2-CPIP. (A–B) CPIP cochaperonin silencing does not interfere with Roq1-triggered hypersensitive response (HR). *Nicotiana benthamiana* plants underwent VIGS with TRV2-GFP (control) or TRV2-CPIP and were then co-infiltrated with Roq1-FLAG, XopQ-STREP, and potato-derived cochaperonins CPIP1-GFP and/or CPIP2a-RFP, as detailed. (A) Shows typical HR phenotypes. (B) Displays immunoblot results verifying protein expression levels.

**Supplementary Figure 2.**
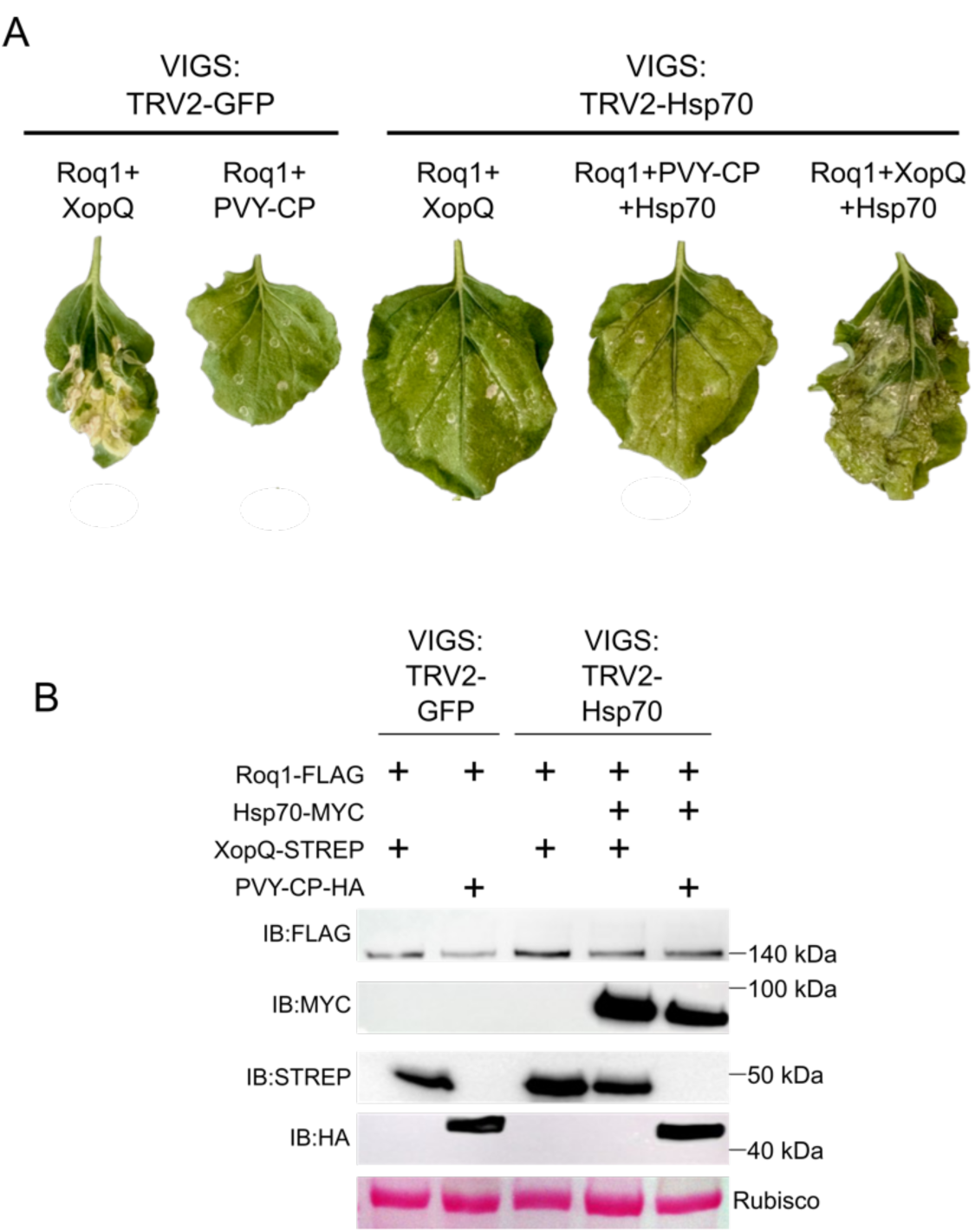
Control analysis of the Roq1–XopQ system in plants silenced for TRV2-GFP or TRV2-Hsp70. (A–B) Hsp70 is essential for Roq1-dependent hypersensitive response (HR) induction. Leaves of *N. benthamiana*, either control TRV2-GFP or TRV2-Hsp70–silenced, were co-infiltrated with Roq1-FLAG and XopQ-STREP, either alone or with potato-derived Hsp70-MYC. PVY CP-HA served as a negative control. (A) Shows typical HR phenotypes. (B) Displays immunoblot results verifying protein levels.

**Supplementary Figure 3.**
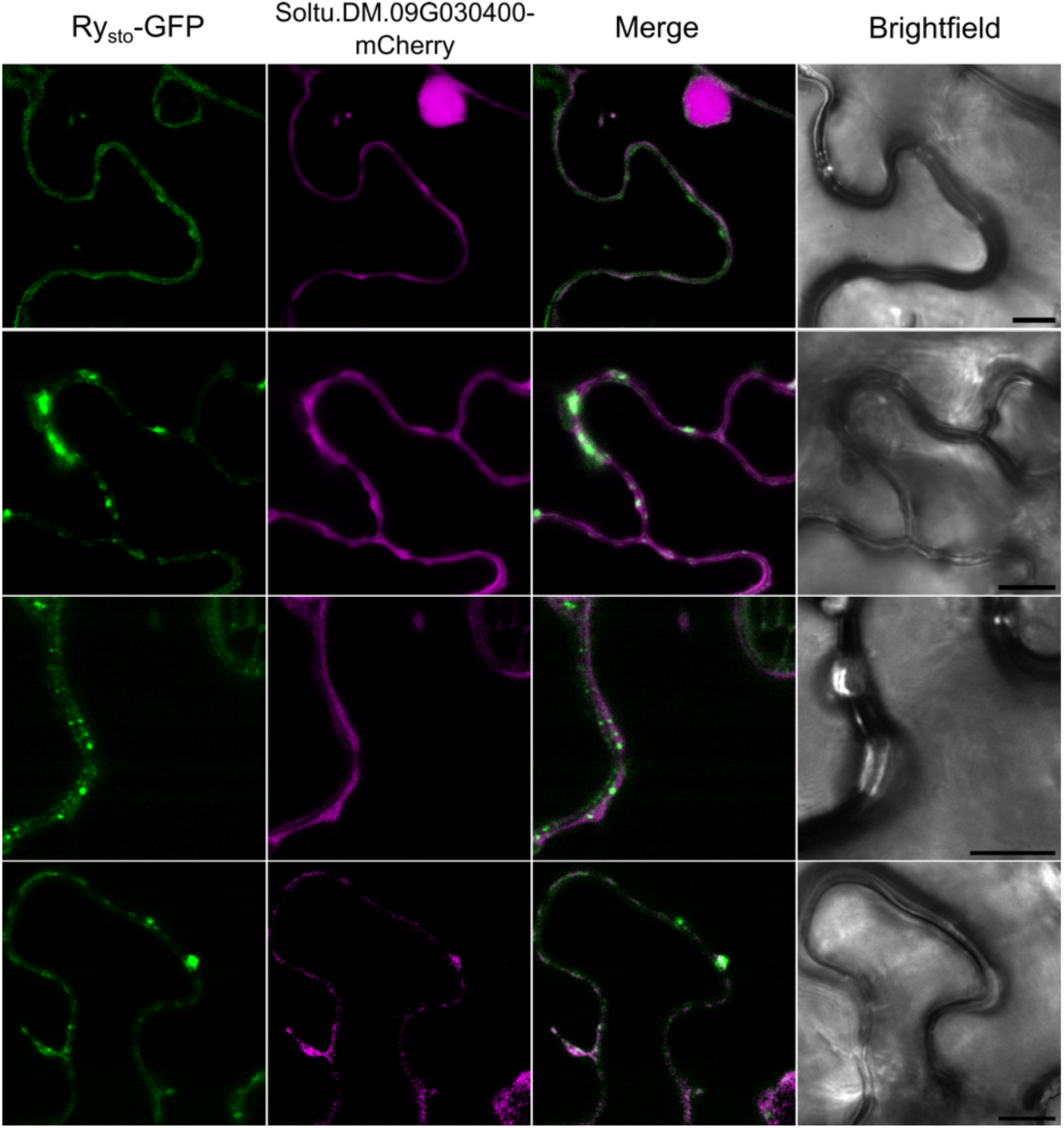
Cortical view of *N. benthamiana* leaf epidermal cells transiently co-expressing Ry_sto_ and free fluorescent proteins (FP). Control plants were co-expressing Ry_sto_-GFP and mCherry. Ry_sto_-RFP was expressed in plants infected with GFP-tagged clones of PVY, TuMV, or PPV. Twenty hours after induction, confocal microscopy was performed to assess the subcellular distribution of Ry_sto_. Scale bars: 10 µm.

**Supplementary Table 1.** Percentage of reads mapped to *S. tuberosum* and PVY genomes. Total and percentage of reads mapped to *S. tuberosum* genome DM_1-3_516_R44 v6.1 and PVY GCF_000862905.1 genomes using HISAT2 and Bowtie2, respectively.

| Genotype | Treatment | Time | Biological repeat | Total reads | Mapped to Stu | Percentage of mapping to Stu | Mapped to PVY | Percentage of mapping to PVY |
| --- | --- | --- | --- | --- | --- | --- | --- | --- |
| WT | Untreated |  | R1 | 86523598 | 74658499 | 86.29% | 24 | 0.00% |
| WT | Untreated |  | R2 | 118447110 | 102613699 | 86.63% | 30 | 0.00% |
| WT | Untreated |  | R3 | 112066048 | 96645716 | 86.24% | 43 | 0.00% |
| *Ry<sub>sto</sub>* | Untreated |  | R1 | 102138924 | 88324792 | 86.48% | 0 | 0.00% |
| *Ry<sub>sto</sub>* | Untreated |  | R2 | 127953510 | 110901719 | 86.67% | 0 | 0.00% |
| *Ry<sub>sto</sub>* | Untreated |  | R3 | 102710210 | 88617359 | 86.28% | 0 | 0.00% |
| WT | mock | 0h | R1 | 85519440 | 73242729 | 85.64% | 23 | 0.00% |
| WT | mock | 0h | R2 | 85256116 | 72791388 | 85.38% | 6 | 0.00% |
| WT | mock | 0h | R3 | 90647544 | 77485472 | 85.48% | 18 | 0.00% |
| *Ry<sub>sto</sub>* | mock | 0h | R1 | 113741466 | 98803593 | 86.87% | 6 | 0.00% |
| *Ry<sub>sto</sub>* | mock | 0h | R2 | 86376810 | 73870117 | 85.52% | 0 | 0.00% |
| *Ry<sub>sto</sub>* | mock | 0h | R3 | 82856842 | 70577534 | 85.18% | 0 | 0.00% |
| WT | mock | 3h | R1 | 85946278 | 72319069 | 84.14% | 0 | 0.00% |
| WT | mock | 3h | R2 | 80686716 | 68111343 | 84.41% | 0 | 0.00% |
| WT | mock | 3h | R3 | 87714182 | 73917048 | 84.27% | 30 | 0.00% |
| *Ry<sub>sto</sub>* | mock | 3h | R1 | 126592880 | 109838574 | 86.77% | 0 | 0.00% |
| *Ry<sub>sto</sub>* | mock | 3h | R2 | 81546980 | 69389682 | 85.09% | 0 | 0.00% |
| *Ry<sub>sto</sub>* | mock | 3h | R3 | 89773786 | 77538602 | 86.37% | 0 | 0.00% |
| WT | PVY | 0h | R1 | 91436002 | 77364413 | 84.61% | 395426 | 0.43% |
| WT | PVY | 0h | R2 | 79935110 | 67370713 | 84.28% | 363418 | 0.45% |
| WT | PVY | 0h | R3 | 101616670 | 85948317 | 84.58% | 444036 | 0.44% |
| *Ry<sub>sto</sub>* | PVY | 0h | R1 | 84605886 | 72145870 | 85.27% | 181143 | 0.21% |
| *Ry<sub>sto</sub>* | PVY | 0h | R2 | 85160028 | 68392106 | 80.31% | 331762 | 0.39% |
| *Ry<sub>sto</sub>* | PVY | 0h | R3 | 84898186 | 72339744 | 85.21% | 316400 | 0.37% |
| WT | PVY | 3h | R1 | 79398576 | 66078394 | 83.22% | 458497 | 0.58% |
| WT | PVY | 3h | R2 | 81737068 | 68199849 | 83.44% | 155267 | 0.19% |
| WT | PVY | 3h | R3 | 83100300 | 69129963 | 83.19% | 367589 | 0.44% |
| *Ry<sub>sto</sub>* | PVY | 3h | R1 | 80769176 | 68719275 | 85.08% | 255836 | 0.32% |
| *Ry<sub>sto</sub>* | PVY | 3h | R2 | 83117578 | 69839950 | 84.03% | 471785 | 0.57% |
| *Ry<sub>sto</sub>* | PVY | 3h | R3 | 88950632 | 74790690 | 84.08% | 409807 | 0.46% |

**Supplementary Table 2.** A List of constructs and organisms used and produced in this study. Organism Strain/Accession *Escherichia coli* DH5α *Agrobacterium tumefaciens* GV3101 *Nicotiana benthamiana eds-1 knockout Nicotiana benhamiana*

| Name | Gene | Tag | Backbone | Promoter | Terminator | Agrobacterium strain | References |
| --- | --- | --- | --- | --- | --- | --- | --- |
| Ry-FLAG | *Ry<sub>sto</sub>* | 3xFLAG-His | pICSL86977OD | 35S | Tocs | GV3101 | This study |
| PVY-CP-HA | *CP_PVY* | HA | pICSL86977OD | 35S | Tocs | GV3101 | This study |
| PVY-CP-FLAG | *CP_PVY* | 3xFLAG | pICSL86977OD | 35S | Tocs | GV3101 | This study |
| CPIP1-GFP | *CPIP1* | GFP | pBAV | pOp6 | NOS | GV3101 | This study |
| CPIP2a-RFP | *CPIP2a* | RFP | pBAV | pOp6 | NOS | GV3101 | This study |
| Roq1-FLAG | *Roq1* | 3xFLAG | pICSL86977OD | 35S | Tocs | GV3101 | This study |
| XopQ-STREP | *XopQ* |  |  |  |  |  | A gift from B. Staskawicz |
| Hsp70-MYC | *StHsp70* | 4xMYC | pGWB | 35s | NOS | GV3101 | This study |
| Soltu.DM.09G030400 | *Soltu.DM.09G030400* | mcherry | pICSL86977OD | 35s | Tocs | Gv3101 | This study |

Organism Strain/Accession
*Escherichia coli* DH5α
*Agrobacterium tumefaciens* GV3101
*Nicotiana benthamiana*
*eds-1* knockout *Nicotiana benthamiana*

