## Supplementary material for "Dynamic Ry_sto_ receptor remodeling controls its ability to confer extreme resistance": dataset 1

| gene_id | gene_name | gene_chr | gene_start | gene_end | gene_strand | gene_length |
|---|---|---|---|---|---|---|
| Soltu.DM.09G | Soltu.DM.09G | chr09 | 6417719 | 6432411 | - | 1920 |
| Soltu.DM.09G | Soltu.DM.09G | chr09 | 59080270 | 59083931 | + | 915 |
| Soltu.DM.09G | Soltu.DM.09G | chr09 | 66722602 | 66724543 | + | 1758 |
| Soltu.DM.09G | Soltu.DM.09G | chr09 | 66346236 | 66347080 | + | 666 |
| Soltu.DM.09G | Soltu.DM.09G | chr09 | 66343871 | 66345138 | + | 833 |
| Soltu.DM.09G | Soltu.DM.09G | chr09 | 66340403 | 66342390 | + | 1415 |
| Soltu.DM.03G | Soltu.DM.03G | chr03 | 2041775 | 2043453 | - | 1496 |
| Soltu.DM.03G | Soltu.DM.03G | chr03 | 6938746 | 6941835 | + | 1228 |
| Soltu.DM.03G | Soltu.DM.03G | chr03 | 59811655 | 59814760 | - | 1462 |
| Soltu.DM.06G | Soltu.DM.06G | chr06 | 42105854 | 42107438 | + | 768 |
| Soltu.DM.06G | Soltu.DM.06G | chr06 | 43156627 | 43159569 | - | 1787 |
| Soltu.DM.06G | Soltu.DM.06G | chr06 | 49856951 | 49858892 | - | 1465 |
| Soltu.DM.06G | Soltu.DM.06G | chr06 | 49851530 | 49852835 | - | 762 |
| Soltu.DM.04G | Soltu.DM.04G | chr04 | 50277361 | 50277753 | + | 393 |
| Soltu.DM.04G | Soltu.DM.04G | chr04 | 3554216 | 3555209 | + | 722 |
| Soltu.DM.04G | Soltu.DM.04G | chr04 | 3571043 | 3571881 | + | 654 |
| Soltu.DM.04G | Soltu.DM.04G | chr04 | 3544234 | 3550053 | + | 1080 |
| Soltu.DM.04G | Soltu.DM.04G | chr04 | 3217164 | 3217992 | - | 668 |
| Soltu.DM.04G | Soltu.DM.04G | chr04 | 2043038 | 2044576 | + | 788 |
| Soltu.DM.04G | Soltu.DM.04G | chr04 | 49203918 | 49207330 | + | 1263 |
| Soltu.DM.04G | Soltu.DM.04G | chr04 | 1470763 | 1474192 | + | 735 |
| Soltu.DM.04G | Soltu.DM.04G | chr04 | 2028744 | 2030217 | + | 709 |
| Soltu.DM.04G | Soltu.DM.04G | chr04 | 1048582 | 1049862 | - | 792 |
| Soltu.DM.04G | Soltu.DM.04G | chr04 | 1464334 | 1468721 | + | 703 |
| Soltu.DM.07G | Soltu.DM.07G | chr07 | 12128096 | 12130223 | + | 927 |
| Soltu.DM.02G | Soltu.DM.02G | chr02 | 34269415 | 34274656 | - | 2005 |
| Soltu.DM.02G | Soltu.DM.02G | chr02 | 27031621 | 27035454 | + | 3000 |
| Soltu.DM.02G | Soltu.DM.02G | chr02 | 17470091 | 17471636 | + | 153 |
| Soltu.DM.11G | Soltu.DM.11G | chr11 | 14416504 | 14418409 | - | 1362 |
| Soltu.DM.08G | Soltu.DM.08G | chr08 | 57933232 | 57934873 | + | 1332 |
| Soltu.DM.08G | Soltu.DM.08G | chr08 | 54564523 | 54568126 | + | 2037 |
| Soltu.DM.08G | Soltu.DM.08G | chr08 | 31646120 | 31649414 | - | 800 |
| Soltu.DM.08G | Soltu.DM.08G | chr08 | 700146 | 709796 | + | 2297 |
| Soltu.DM.08G | Soltu.DM.08G | chr08 | 6462121 | 6462886 | - | 672 |
| Soltu.DM.08G | Soltu.DM.08G | chr08 | 40434955 | 40439383 | - | 673 |
| Soltu.DM.08G | Soltu.DM.08G | chr08 | 43874268 | 43876854 | + | 1616 |
| Soltu.DM.08G | Soltu.DM.08G | chr08 | 48490878 | 48492116 | + | 1239 |
| Soltu.DM.10G | Soltu.DM.10G | chr10 | 52203586 | 52204550 | - | 965 |
| Soltu.DM.10G | Soltu.DM.10G | chr10 | 15165640 | 15167637 | + | 822 |
| Soltu.DM.05G | Soltu.DM.05G | chr05 | 31325385 | 31326282 | - | 498 |
| Soltu.DM.05G | Soltu.DM.05G | chr05 | 9840320 | 9842588 | - | 1683 |
| Soltu.DM.12G | Soltu.DM.12G | chr12 | 57834518 | 57837110 | - | 957 |
| Soltu.DM.12G | Soltu.DM.12G | chr12 | 16113176 | 16120802 | + | 645 |
| Soltu.DM.12G | Soltu.DM.12G | chr12 | 10977629 | 10979305 | + | 1339 |
| Soltu.DM.12G | Soltu.DM.12G | chr12 | 50992224 | 50995223 | - | 1687 |
| Soltu.DM.12G | Soltu.DM.12G | chr12 | 49161389 | 49163549 | - | 330 |
| Soltu.DM.12G | Soltu.DM.12G | chr12 | 2713161 | 2717150 | + | 2371 |
| Soltu.DM.01G | Soltu.DM.01G | chr01 | 69393561 | 69397985 | + | 2014 |
| Soltu.DM.01G | Soltu.DM.01G | chr01 | 77724204 | 77728082 | + | 2800 |

| | | | | | | |
|---|---|---|---|---|---|---|
| Soltu.DM.01G | Soltu.DM.01G | chr01 | 88209323 | 88211644 | - | 1708 |
| Soltu.DM.01G | Soltu.DM.01G | chr01 | 29940213 | 29942321 | + | 867 |
| Soltu.DM.01G | Soltu.DM.01G | chr01 | 41858204 | 41860939 | - | 423 |
| Soltu.DM.01G | Soltu.DM.01G | chr01 | 72727897 | 72728787 | - | 891 |
| Soltu.DM.01G | Soltu.DM.01G | chr01 | 85537996 | 85539146 | - | 874 |
| Soltu.DM.01G | Soltu.DM.01G | chr01 | 72773873 | 72776361 | - | 1749 |
| Soltu.DM.01G | Soltu.DM.01G | chr01 | 82995620 | 82998696 | + | 1830 |
| Soltu.DM.01G | Soltu.DM.01G | chr01 | 82625019 | 82626820 | - | 507 |
| Soltu.DM.01G | Soltu.DM.01G | chr01 | 72729916 | 72732827 | - | 2181 |
| Soltu.DM.01G | Soltu.DM.01G | chr01 | 15933649 | 15934904 | - | 687 |
| Soltu.DM.01G | Soltu.DM.01G | chr01 | 78376335 | 78378723 | - | 1376 |
| Soltu.DM.01G | Soltu.DM.01G | chr01 | 65982650 | 65983715 | + | 949 |
| Soltu.DM.01G | Soltu.DM.01G | chr01 | 72792945 | 72794363 | - | 1342 |

| gene_biotype | gene_descript | Family |
| --- | --- | --- |
| protein_codin | - && sp\|Q6NL | ERF |
| protein_codin | - && - && - | - |
| protein_codin | - && sp\|Q8VV | MYB |
| protein_codin | - && - && - | - |
| protein_codin | - && - && - | - |
| protein_codin | - && - && - | - |
| protein_codin | - && sp\|Q84V | - |
| protein_codin | - && sp\|W8JV | GRF |
| protein_codin | - && sp\|Q9LJI | C3H |
| protein_codin | - && sp\|Q9FC | GRAS |
| protein_codin | - && sp\|Q9FC | GRAS |
| protein_codin | - && sp\|Q945 | Dof |
| protein_codin | - && sp\|Q945 | Dof |
| protein_codin | - && sp\|Q940 | - |
| protein_codin | - && sp\|P855 | - |
| protein_codin | - && sp\|P855 | - |
| protein_codin | - && sp\|P855 | - |
| protein_codin | - && sp\|P855 | - |
| protein_codin | - && sp\|Q941 | - |
| protein_codin | - && sp\|Q9XE | MYB |
| protein_codin | - && sp\|P855 | - |
| protein_codin | - && sp\|P855 | - |
| protein_codin | - && sp\|P855 | - |
| protein_codin | - && sp\|P855 | - |
| protein_codin | - && - && PFC | Trihelix |
| protein_codin | - && sp\|Q9LV | MYB_related |
| protein_codin | - && sp\|Q9SH | GATA |
| protein_codin | - && - && - | - |
| protein_codin | - && sp\|Q3ZP | ERF |
| protein_codin | - && sp\|Q9FP | - |
| protein_codin | - && sp\|O652 | - |
| protein_codin | - && sp\|P198 | - |
| protein_codin | - && sp\|Q93Z | C3H |
| protein_codin | - && - && PFC | - |
| protein_codin | - && - && PFC | - |
| protein_codin | - && sp\|Q9SA | WRKY |
| protein_codin | - && sp\|P142 | bZIP |
| protein_codin | - && - && - | - |
| protein_codin | - && - && - | - |
| protein_codin | - && sp\|Q0W | CAMTA |
| protein_codin | - && sp\|Q9LP | - |
| protein_codin | - && - && PFC | bHLH |
| protein_codin | - && - && - | NF-YA |
| protein_codin | - && - && PF1 | WRKY |
| protein_codin | - && sp\|Q9FC | GRAS |
| protein_codin | - && - && - | B3 |
| protein_codin | - && sp\|Q403 | B3 |
| protein_codin | - && sp\|B8BH | - |
| protein_codin | - && sp\|Q431 | NF-YA |

| | | |
|---|---|---|
| protein_codin | - && sp\|Q9LM | C2H2 |
| protein_codin | - && - && PF0 | - |
| protein_codin | - && - && - | - |
| protein_codin | - && sp\|Q9FZ | ERF |
| protein_codin | - && sp\|Q9SE | NF-YB |
| protein_codin | - && sp\|Q50E | NAC |
| protein_codin | - && sp\|G1JU | bHLH |
| protein_codin | - && - && - | - |
| protein_codin | - && sp\|Q50E | NAC |
| protein_codin | - && sp\|P855 | - |
| protein_codin | - && sp\|O804 | GRAS |
| protein_codin | - && sp\|Q6K5 | E2F/DP |
| protein_codin | - && sp\|O238 | - |
