## Supplementary material for "Dynamic Ry_sto_ receptor remodeling controls its ability to confer extreme resistance": dataset 2

| Gene ID | Size | Database | Predicted domain |
|---|---|---|---|
| Soltu.DM.09G030380.1 | 448 | Phobius | NON_CYTOPLASMIC_DOMAIN |
| Soltu.DM.09G030380.1 | 448 | SignalP_GRAM_POSITIVE | SignalP-TM |
| Soltu.DM.09G030380.1 | 448 | Phobius | SIGNAL_PEPTIDE |
| Soltu.DM.09G030380.1 | 448 | SignalP_EUK | SignalP-noTM |
| Soltu.DM.09G030380.1 | 448 | Phobius | SIGNAL_PEPTIDE_C_REGION |
| Soltu.DM.09G030380.1 | 448 | Phobius | SIGNAL_PEPTIDE_H_REGION |
| Soltu.DM.09G030380.1 | 448 | Phobius | SIGNAL_PEPTIDE_N_REGION |
| Soltu.DM.09G030380.1 | 448 | MobiDBLite | mobidb-lite |
| Soltu.DM.09G030400.1 | 100 | PANTHER | PTHR37389 |
| Soltu.DM.09G030400.1 | 100 | SignalP_GRAM_POSITIVE | SignalP-TM |
| Soltu.DM.09G030400.1 | 100 | TMHMM | TMhelix |
| Soltu.DM.09G030400.1 | 100 | SignalP_EUK | SignalP-noTM |
| Soltu.DM.09G030400.1 | 100 | Phobius | SIGNAL_PEPTIDE |
| Soltu.DM.09G030400.1 | 100 | Phobius | SIGNAL_PEPTIDE_C_REGION |
| Soltu.DM.09G030400.1 | 100 | Phobius | SIGNAL_PEPTIDE_H_REGION |
| Soltu.DM.09G030400.1 | 100 | Phobius | SIGNAL_PEPTIDE_N_REGION |
| Soltu.DM.09G030400.1 | 100 | Phobius | NON_CYTOPLASMIC_DOMAIN |
| Soltu.DM.09G030390.1 | 118 | TMHMM | TMhelix |
| Soltu.DM.09G030390.1 | 118 | PANTHER | PTHR37389 |
| Soltu.DM.09G030390.1 | 118 | TMHMM | TMhelix |
| Soltu.DM.09G030390.1 | 118 | SignalP_EUK | SignalP-noTM |
| Soltu.DM.09G030390.1 | 118 | SignalP_GRAM_POSITIVE | SignalP-TM |
| Soltu.DM.09G030390.1 | 118 | Phobius | NON_CYTOPLASMIC_DOMAIN |
| Soltu.DM.09G030390.1 | 118 | Phobius | SIGNAL_PEPTIDE |
| Soltu.DM.09G030390.1 | 118 | Phobius | SIGNAL_PEPTIDE_C_REGION |
| Soltu.DM.09G030390.1 | 118 | Phobius | SIGNAL_PEPTIDE_H_REGION |
| Soltu.DM.09G030390.1 | 118 | Phobius | SIGNAL_PEPTIDE_N_REGION |
| Soltu.DM.09G030390.1 | 118 | TMHMM | TMhelix |
| Soltu.DM.09G030410.1 | 161 | SignalP_EUK | SignalP-noTM |
| Soltu.DM.09G030410.1 | 161 | Phobius | NON_CYTOPLASMIC_DOMAIN |
| Soltu.DM.09G030410.1 | 161 | SignalP_GRAM_POSITIVE | SignalP-TM |
| Soltu.DM.09G030410.1 | 161 | Phobius | SIGNAL_PEPTIDE |
| Soltu.DM.09G030410.1 | 161 | PANTHER | PTHR37389 |
| Soltu.DM.09G030410.1 | 161 | TMHMM | TMhelix |
| Soltu.DM.09G030410.1 | 161 | Phobius | SIGNAL_PEPTIDE_C_REGION |
| Soltu.DM.09G030410.1 | 161 | Phobius | SIGNAL_PEPTIDE_H_REGION |
| Soltu.DM.09G030410.1 | 161 | Phobius | SIGNAL_PEPTIDE_N_REGION |
| Soltu.DM.09G030370.1 | 208 | Phobius | NON_CYTOPLASMIC_DOMAIN |
| Soltu.DM.09G030370.1 | 208 | TMHMM | TMhelix |
| Soltu.DM.09G030370.1 | 208 | Phobius | SIGNAL_PEPTIDE |
| Soltu.DM.09G030370.1 | 208 | TMHMM | TMhelix |
| Soltu.DM.09G030370.1 | 208 | SignalP_GRAM_POSITIVE | SignalP-TM |
| Soltu.DM.09G030370.1 | 208 | Phobius | SIGNAL_PEPTIDE_C_REGION |

| | | | |
|---|---|---|---|
| Soltu.DM.09G030370.1 | 208 | Phobius | SIGNAL_PEPTIDE_H_REGION |
| Soltu.DM.09G030370.1 | 208 | Phobius | SIGNAL_PEPTIDE_N_REGION |
| Soltu.DM.09G030370.1 | 208 | SignalP_EUK | SignalP-noTM |

| Description |
|---|
| Region of a membrane-bound protein predicted to be outside the membrane, in the extracellular region. |
| SignalP-TM |
| Signal peptide region |
| SignalP-noTM |
| C-terminal region of a signal peptide. |
| Hydrophobic region of a signal peptide. |
| N-terminal region of a signal peptide. |
| consensus disorder prediction |
| NODULIN-24 |
| SignalP-TM |
| Region of a membrane-bound protein predicted to be embedded in the membrane. |
| SignalP-noTM |
| Signal peptide region |
| C-terminal region of a signal peptide. |
| Hydrophobic region of a signal peptide. |
| N-terminal region of a signal peptide. |
| Region of a membrane-bound protein predicted to be outside the membrane, in the extracellular region. |
| Region of a membrane-bound protein predicted to be embedded in the membrane. |
| NODULIN-24 |
| Region of a membrane-bound protein predicted to be embedded in the membrane. |
| SignalP-noTM |
| SignalP-TM |
| Region of a membrane-bound protein predicted to be outside the membrane, in the extracellular region. |
| Signal peptide region |
| C-terminal region of a signal peptide. |
| Hydrophobic region of a signal peptide. |
| N-terminal region of a signal peptide. |
| Region of a membrane-bound protein predicted to be embedded in the membrane. |
| SignalP-noTM |
| Region of a membrane-bound protein predicted to be outside the membrane, in the extracellular region. |
| SignalP-TM |
| Signal peptide region |
| NODULIN-24 |
| Region of a membrane-bound protein predicted to be embedded in the membrane. |
| C-terminal region of a signal peptide. |
| Hydrophobic region of a signal peptide. |
| N-terminal region of a signal peptide. |
| Region of a membrane-bound protein predicted to be outside the membrane, in the extracellular region. |
| Region of a membrane-bound protein predicted to be embedded in the membrane. |
| Signal peptide region |
| Region of a membrane-bound protein predicted to be embedded in the membrane. |
| SignalP-TM |
| C-terminal region of a signal peptide. |

| |
|---|
| Hydrophobic region of a signal peptide. |
| N-terminal region of a signal peptide. |
| SignalP-noTM |

| Start | End | Column1 | Column2 | Protein Family |
|---|---|---|---|---|
| 24 | 448 | - | - | - |
| 1 | 22 | - | - | - |
| 1 | 23 | - | - | - |
| 1 | 23 | - | - | - |
| 17 | 23 | - | - | - |
| 5 | 16 | - | - | - |
| 1 | 4 | - | - | - |
| 291 | 372 | - | - | - |
| 1 | 72 | 3,20E-10 | IPR010800 | Glycine rich protein |
| 1 | 22 | - | - | - |
| 5 | 22 | - | - | - |
| 1 | 23 | - | - | - |
| 1 | 23 | - | - | - |
| 17 | 23 | - | - | - |
| 5 | 16 | - | - | - |
| 1 | 4 | - | - | - |
| 24 | 100 | - | - | - |
| 95 | 117 | - | - | - |
| 1 | 103 | 1,50E-14 | IPR010800 | Glycine rich protein |
| 5 | 27 | - | - | - |
| 1 | 23 | - | - | - |
| 1 | 22 | - | - | - |
| 24 | 118 | - | - | - |
| 1 | 23 | - | - | - |
| 17 | 23 | - | - | - |
| 5 | 16 | - | - | - |
| 1 | 4 | - | - | - |
| 48 | 70 | - | - | - |
| 1 | 23 | - | - | - |
| 24 | 161 | - | - | - |
| 1 | 22 | - | - | - |
| 1 | 23 | - | - | - |
| 1 | 107 | 1,30E-13 | IPR010800 | Glycine rich protein |
| 5 | 23 | - | - | - |
| 19 | 23 | - | - | - |
| 8 | 18 | - | - | - |
| 1 | 7 | - | - | - |
| 24 | 208 | - | - | - |
| 5 | 27 | - | - | - |
| 1 | 23 | - | - | - |
| 47 | 69 | - | - | - |
| 1 | 22 | - | - | - |
| 17 | 23 | - | - | - |

| | | | | |
|---|---|---|---|---|
| 5 | 16 | - | - | - |
| 1 | 4 | - | - | - |
| 1 | 23 | - | - | - |
